# CNS-PENETRANT NLRP3 INHIBITOR ACHIEVES DURABLE WEIGHT LOSS AND REVERSES HYPOTHALAMIC INFLAMMATION IN DIET-INDUCED OBESITY

**DOI:** 10.1101/2025.10.06.680810

**Authors:** Jing Guo, Tomas P. Bachor, Ryan Takahashi, Micah Steffek, Gabriel A. Fitzgerald, Benjamin J. Huffman, Dylan Braun, Suk Ji Han, Thomas Sandmann, Connie Ha, Katrina W. Lexa, Yun-Ting Hsieh, Maayan R. Agam, Elizabeth W. Sun, Xiang Wang, Min Liu, Javier de Vicente, Brian M. Fox, Meghan K. Campbell, Stephen P. Cavnar, Tracy J. Yuen, Tanya Z. Fischer, Anthony A. Estrada, Robert A. Craig, Ernie Yulyaningsih

**Author notes:** Corresponding author: Ernie Yulyaningsih, PhD Tenvie Therapeutics, 1400 Sierra Point Parkway, Suite 300, Brisbane, CA 94005.

## Abstract

The NLRP3 inflammasome is a key mediator of innate immunity that integrates inflammatory and metabolic stress signals. Increased and/or chronic activation of this critical pathway has been implicated in obesity, with hypothalamic neuroinflammation linked to dysregulation of energy balance. TN-783 is an investigational, CNS-penetrant, small-molecule NLRP3 inhibitor that potently suppressed inflammasome activation across multiple *in vitro* assays. In diet-induced obese (DIO) mice, only TN-783, and not the peripherally restricted NLRP3 inhibitor TN-101, produced progressive and sustained weight loss, underscoring the requirement for central target engagement. Weight loss was driven by a persistent reduction in food intake across both acute and chronic phases, without altering energy expenditure. This effect was further characterized by selective reduction of fat mass, with minimal impact on lean tissues. Mechanistically, NLRP3 inhibition attenuated DIO-induced hypothalamic neuroinflammation and partially reversed obesity-associated molecular changes based on transcriptomic and proteomic profiling of the hypothalamus. Beyond monotherapy, TN-783 enhanced the effects of the GLP-1 receptor agonist semaglutide by amplifying weight loss, reinitiating weight loss after semaglutide effect had plateaued, and maintaining the weight loss benefit after semaglutide withdrawal. Discontinuation of TN-783 resulted in reversal of both weight and feeding effects, indicating that its therapeutic activity requires ongoing target engagement rather than permanent remodeling of metabolic pathways. Collectively, these observations support central NLRP3 inhibition as a distinct and promising approach for obesity treatment, offering robust induction and sustained maintenance of weight loss while preserving reversibility.

## INTRODUCTION

Obesity is a chronic, multifactorial disease shaped by genetic, environmental, and neurobiological influences. While lifestyle modification offers measurable health benefits, durable weight reduction remains difficult for most individuals, and obesity continues to drive rising rates of type 2 diabetes, cardiovascular disease, and other metabolic complications. Pharmacologic treatment has advanced with glucagon-like peptide-1 receptor agonists (GLP-1RA), which promote substantial weight loss through appetite suppression and improved metabolic regulation (1–3). Other incretin-based agents and newer classes of therapies, such as those targeting PYY and amylin, are also emerging (4–7). However, important limitations remain: (1) responses are heterogeneous with many patients failing to achieve a clinically meaningful weight loss, (2) the loss of lean mass can occur during treatment, (3) tolerability issues can hinder adherence, and (4) weight regain often follows treatment withdrawal (8, 9). These challenges underscore the need for complementary strategies that engage distinct mechanisms to improve induction, durability, and quality of weight loss.

Obesity is recognized as a state of chronic, low-grade inflammation, with consistent evidence of elevated circulating inflammatory mediators in affected individuals (10–15). Growing evidence further implicates a central role for neuroinflammation in the pathophysiology of obesity; in preclinical models, high fat feeding and chronic overnutrition trigger activation of microglia and astrocytes in hypothalamic nuclei that regulate energy balance, leading to impaired neuronal signaling and sustained hyperphagia (16–19). Consistent with these preclinical findings, post-mortem analyses of brains from obese patients have revealed microgliosis and exacerbated microglial dystrophy in corresponding hypothalamic regions, further supporting neuroinflammation as a common, driving feature of obesity in both preclinical models and humans (20).

The NOD-, LRR- and pyrin domain-containing protein 3 (NLRP3) inflammasome has been implicated as a mechanistic driver of obesity-associated neuroinflammation and metabolic dysfunction. Activation of NLRP3 induces the assembly of the inflammasome complex with the apoptosis-associated speck-like protein containing a CARD (ASC), leading to caspase-1 activation and the subsequent release of proinflammatory cytokines, IL-1β and IL-18. Mice lacking NLRP3 or downstream components such as ASC or caspase-1 demonstrated attenuated diet-induced weight gain, adipose inflammation, and insulin resistance (21, 22). Additionally, pharmacological studies showed that small molecule NLRP3 inhibitors reduced obesity, attenuated inflammatory biomarkers, and improved metabolic outcomes in rodent models (23, 24). Collectively, these data suggest that the NLRP3 pathway may be a key contributor to the pathophysiology of obesity and thus represents a promising therapeutic target.

In this study, we developed two structurally distinct NLRP3 inhibitors with differential brain penetrance: TN-783, a highly CNS-penetrant compound, and TN-101, a peripherally restricted analog, to dissect the contribution of central *versus* peripheral NLRP3 inflammasome signaling in obesity. Using DIO mice, we demonstrated that central NLRP3 inhibition was required to produce weight loss, and that this was achieved through selective reduction of fat mass while preserving lean mass. This effect was sustained by a persistent suppression of food intake without compensatory reduction in energy expenditure, across both the acute and chronic phases of weight loss. Beyond amplifying the initial response to semaglutide, NLRP3 inhibition also reinitiated weight loss after GLP-1RA-induced weight loss had plateaued. Mechanistically, TN-783 reduced DIO-induced neuroinflammation in the hypothalamic arcuate nucleus (ARC) and elicited partial restoration of obesity-associated transcriptomic and proteomic dysregulation, most notably within inflammatory and neuronal signaling pathways. Collectively, these findings established the mechanistic and pharmacologic requirements for NLRP3-targeted therapies, resolved uncertainties regarding the necessity of CNS engagement, and demonstrated the potential of NLRP3 inhibition to build upon the benefits of incretin-based treatments.

## RESULTS

### Discovery and characterization of highly CNS-penetrant NLRP3 inhibitors

To identify CNS-penetrant NLRP3 small molecules, we initiated a lead-finding campaign utilizing a ligand-based structure-activity relationship strategy in parallel to a high-throughput screening campaign. Through our lead-finding efforts, we identified TN-551, a compound that showed high potency for NLRP3 when tested in a fluorescence polarization (FP) displacement assay using a fluorescent analog of MCC950 – a well-characterized, sulfonylurea-based reference inhibitor of the NLRP3 inflammasome (25).

To confirm the mechanism of action of this early lead, we determined the cryo-electron microscopy (cryo-EM) structure of the NLRP3–NEK7 complex in the presence of ADP, Mg^2+^, complexed with TN-551 (**Supplementary Figure 1**). This structure provided the basis for iterative rounds of structure-based drug design to improve *in vitro* assay potency and pharmacokinetic properties resulting in the development of TN-783 and TN-101 (**Figure 1A-B**). In our FP displacement assay, TN-101 and TN-783 yielded IC_50_ values of 29.8 ± 4.9 nM and 19.3 ± 2.2 nM (**Figure 1C**, **Supplementary Table 2**), respectively. These data confirm that both compounds bind to the NLRP3 NACHT pocket with high affinity.

**Figure 1.**
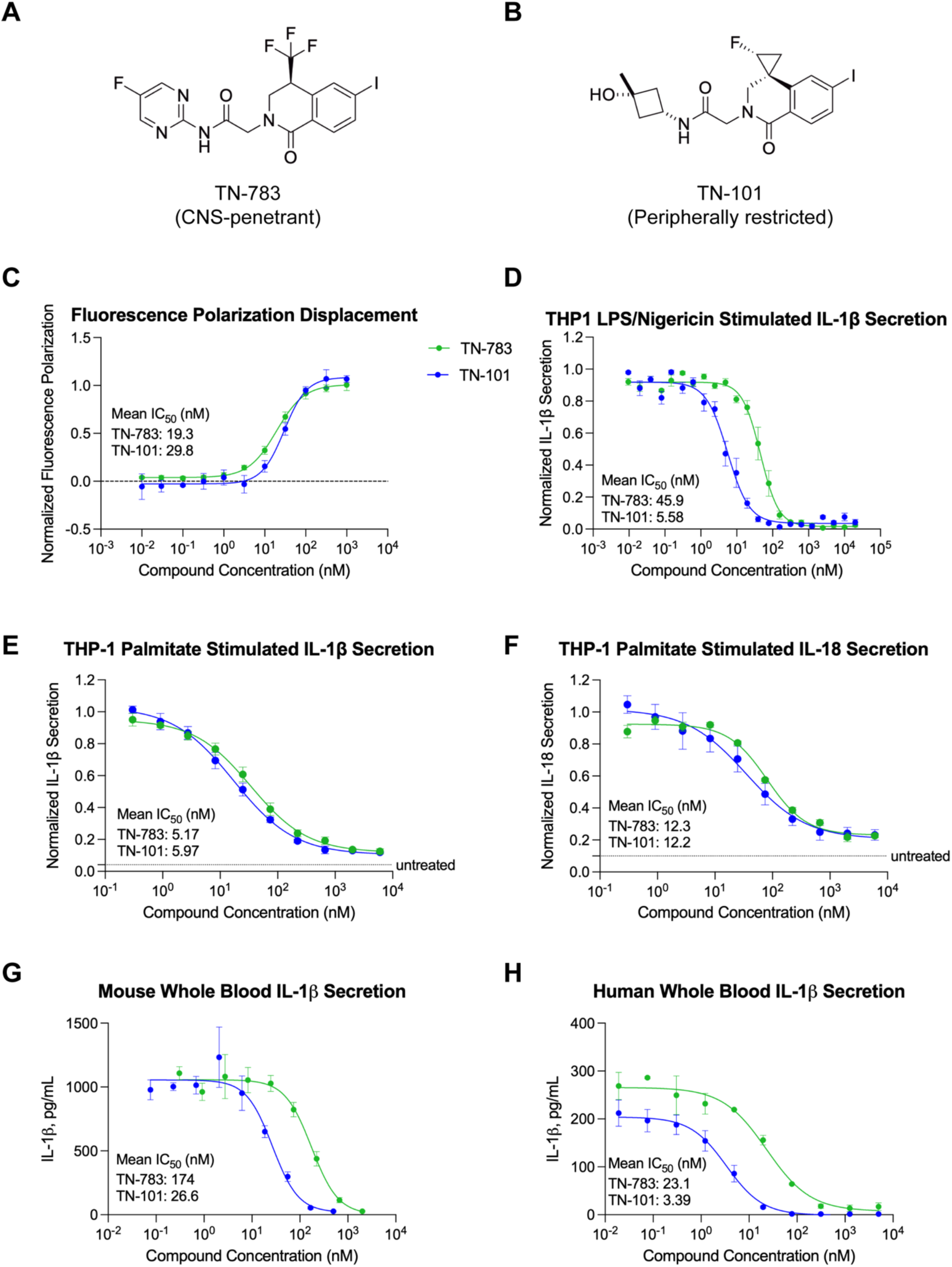
*In vitro* characterization of peripherally restricted and CNS-penetrant NLRP3 inhibitors. (**A-B**) Chemical structures of TN-783 (CNS-penetrant NLRP3 inhibitor) and TN-101 (peripherally restricted NLRP3 inhibitor). (**C**) Dose response curve of TN-101 and TN-783 in fluorescence polarization displacement assay. (**D**) Dose-response curves of TN-101 and TN-783 on IL-1β secretion from differentiated THP-1 macrophages following LPS/nigericin stimulation. Data were normalized by in-plate DMSO-treated and positive control compound-treated conditions. (**E-F**) Dose-response curves of TN-101 and TN-783 on IL-1β (**E**) and IL-18 (**F**) secretion from differentiated THP-1 macrophages following overnight stimulation with 400 μM palmitate-BSA. Concentration of IL-1β or IL-18 detected in the medium at each compound concentration was normalized to the value observed with DMSO control. The dotted line represents the level of IL-1β or IL-18 detected without palmitate treatment (untreated). IC_50_ values were corrected for palmitate-BSA binding of the compounds. (**G-H**) Dose-response curves of TN-101 and TN-783 on IL-1β secretion from ATP/LPS stimulated mouse whole blood (**G**) or human whole blood (**H**). For figures C-H, data are shown as mean ± SEM from three independent experiments.

To characterize the functional effects of our NLRP3 inhibitors, we assessed TN-783 and TN-101 in cellular assays with LPS/nigericin-stimulated THP-1 macrophages, measuring IL-1β release as a readout of inflammasome activation (26). TN-101 and TN-783 inhibited IL-1β secretion with an IC_50_ of 5.58 ± 0.47 nM and 45.9 ± 3.0 nM, respectively (**Figure 1D**, **Supplementary Table 2**), demonstrating strong cellular activity for both compounds in conditions known to stimulate NLRP3-dependent immune responses.

Palmitate, a dietary saturated fatty acid (SFA), is known to induce NLRP3 inflammasome activation, leading to impaired insulin signaling (27, 28). Elevated levels of free fatty acids, including palmitate, have been reported in both the serum and cerebrospinal fluid of obese individuals (29–31). In line with these findings, we confirmed that lipid challenge in THP-1 macrophages with palmitate-BSA robustly stimulated the secretion of IL-1β and IL-18, the key cytokines that are dependent on NLRP3 pathway activation (**Figure 1E-F**). Notably, both NLRP3 inhibitors, TN-101 and TN-783, dose dependently suppressed palmitate-induced inflammatory response, with IC_50_ values of 5.97 ± 0.94 nM and 5.17 ± 0.73 nM for IL-1β, and 12.2 ± 4.3 nM and 12.3 ± 1.4 nM for IL-18 secretion, respectively (**Supplementary Table 2**).

Finally, to determine whether our inhibitors can inhibit NLRP3 activity in translationally relevant samples, we measured IL-1β secretion in mouse whole blood stimulated with LPS/ATP. TN-101 and TN-783 inhibited IL-1β release with IC_50_ values of 26.6 ± 7.1 nM and 174 ± 37 nM, respectively (**Figure 1G)**. In human whole blood, both compounds also showed robust activity, with IC_50_ values of 3.39 ± 0.83 nM for TN-101 and 23.1 ± 6.0 nM for TN-783 (**Figure 1H**). Together, these data demonstrate that TN-783 and TN-101 have comparable efficacy and potency across biophysical and cellular assays.

### Pharmacokinetic analysis demonstrated that TN-783 is highly CNS-penetrant while TN-101 is peripherally restricted

To evaluate the pharmacokinetic (PK) profiles of TN-783 and TN-101 *in vivo*, we conducted dosing studies in both normal chow-fed lean (NC lean) and DIO mice. TN-783 was administered twice daily (*b.i.d.*) *via* oral gavage at 50 mg/kg for 14 days (**Figure 2A**). At steady state, plasma concentrations reached similar peak levels in NC lean and DIO mice, but declined more slowly in the DIO animals, consistent with sustained or slower oral absorption (**Figure 2B**). Under the *b.i.d.* regimen with 10 hours between daily doses, plasma exposures of TN-783 were broadly maintained during the 0-10- and 10-20-hour intervals, with a more pronounced decline occurring between 20-24 hours before the next day’s dose (**Figure 2B**).

**Figure 2.**
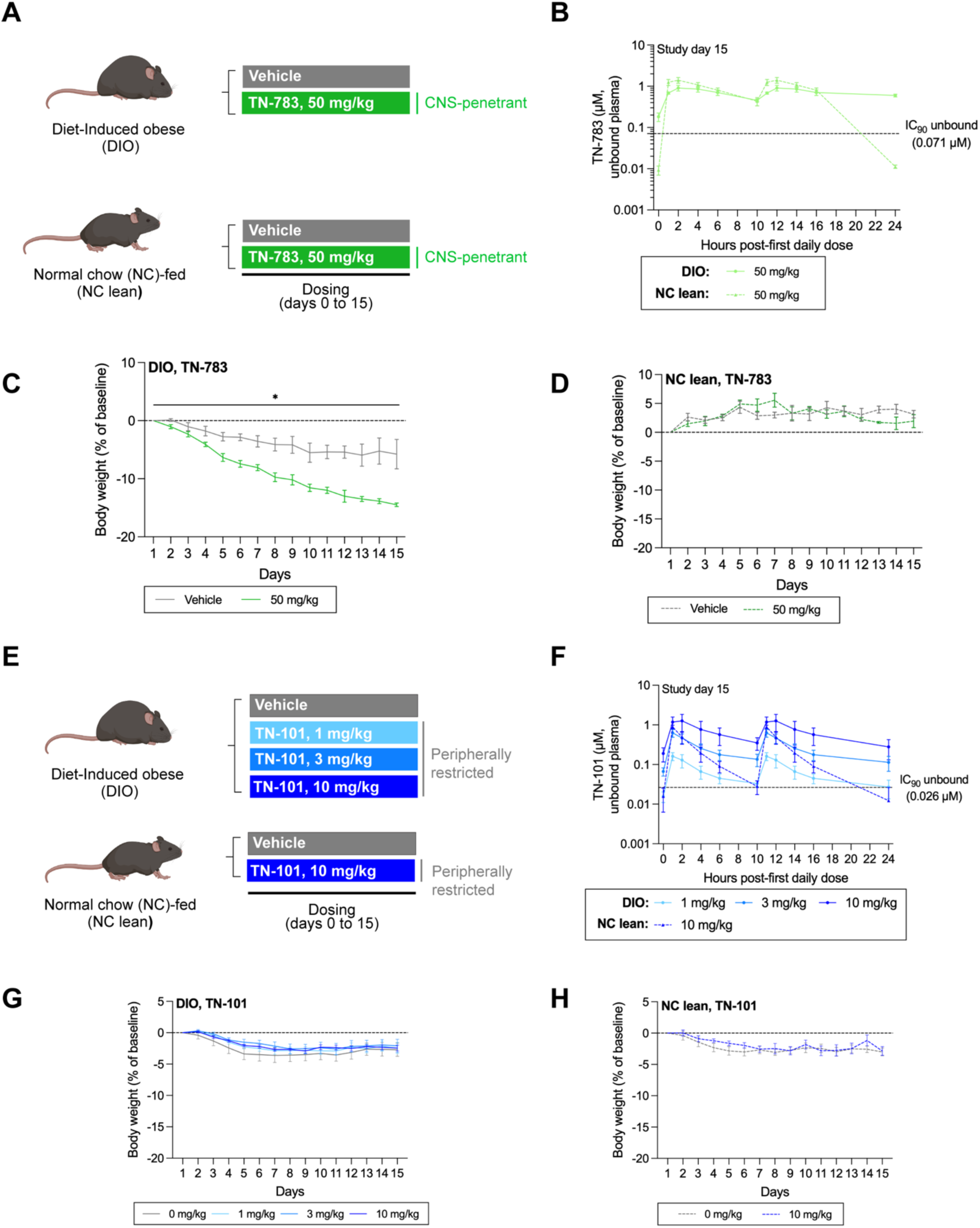
*In vivo* exposure of CNS-penetrant and peripherally restricted NLRP3 inhibitors in DIO and lean mice. (**A**) Schematic of TN-783 dosing in DIO and normal chow (NC)-fed lean mice. Mice received twice daily oral dosing of vehicle or TN-783 (50 mg/kg) for 15 days. (**B**) Unbound plasma TN-783 exposure at steady state (day 15). (**C-D**) Body weight changes from baseline in DIO and NC lean animals. (**E**) Schematic of TN-101 dosing study in DIO and NC lean mice. TN-101 or its vehicle was administered twice daily *via* oral gavage for 15 days. (**F**) Unbound plasma TN-101 exposure at steady state (day 15). (**G-H**) Body weight changes from baseline in DIO and NC lean animals. Data are mean ± SEM of N=3/group (**A-D**) or N=10/group (**E-H**). Statistical significance relative to vehicle-treated control was determined by a two-way ANOVA (*, p<0.05).

Unbound fractions (fu) determined by ultracentrifugation in rodent matrices were 0.045 in plasma and 0.043 in brain for TN-783. To calculate unbound brain exposure, unbound plasma concentrations were scaled using a brain-to-plasma unbound partition ratio (K_p,uu_) of 0.41, derived from dedicated PK studies in DIO mice (**Supplementary Table 3**). At 50 mg/kg, unbound plasma exposures exceeded the *in vitro* unbound IC_90_ (IC_90,u_) for NLRP3 inhibition and calculated unbound brain concentrations also surpassed this benchmark (**Supplementary Table 3**). Administration of TN-783 at 50 mg/kg selectively reduced body weight in DIO mice, whereas lean animals were unaffected (**Figure 2C-D**). This result, while obtained in PK-dedicated cohorts with small group sizes (N=3/group), indicated the weight-lowering effect of a CNS-penetrant NLRP3 inhibitor may be specific to the obese state.

To directly contrast the CNS-penetrant compound with a peripherally restricted analog, we performed parallel studies with TN-101 (**Figure 2E–H**). Like TN-783, TN-101 achieved comparable peak plasma concentrations in NC lean and DIO mice, with exposures again declining more rapidly in NC lean animals. TN-101 displayed approximately dose-proportional increases in steady-state plasma exposure across the dose range of 1 to 10 mg/kg (**Figure 2F**). Unbound fractions in rodent matrices were 0.110 in plasma and 0.144 in brain. Predicted brain concentrations were then derived from unbound plasma exposures using a K_p,uu_ of 0.016, established in dedicated PK studies in DIO mice (**Supplementary Table 3**). Under these conditions, unbound plasma exposures exceeded the *in vitro* unbound IC_90_ (IC_90,u_) for NLRP3 inhibition at all doses while unbound brain levels remained below the *in vitro* IC_90,u_ for NLRP3 inhibition, even at the highest dose. Thus, while TN-101 reached adequate systemic exposure, CNS exposure was comparatively restricted. Associated with this, TN-101 did not induce body weight change in DIO mice at all doses tested up to 10 mg/kg (**Figure 2G**). Treatment of lean mice with the highest dose of TN-101 for 15 days was well tolerated and did not alter body weight relative to vehicle controls (**Figure 2H**), suggesting the absence of weight change in DIO mice is not due to or confounded by adverse events. On this basis, 1 mg/kg was selected for future studies, as it achieved plasma exposure above IC_90,u_ while maintaining minimal CNS penetration.

Taken together, the body weight reduction induced by the CNS-penetrant NLRP3 inhibitor TN-783 and the absence of body weight change after dosing with the peripherally restricted NLRP3 inhibitor, TN-101, is consistent with the hypothesis that CNS NLRP3 pathway inhibition is required for efficacy.

### CNS-penetrant NLRP3 inhibitor promoted progressive and sustained weight loss in DIO mice

Inhibition of NLRP3 has been previously shown to induce weight loss in models of obesity (23, 24), but whether this effect was mediated by central or peripheral inflammasome activity remained unresolved. Our PK study provided an early indication that TN-783, a CNS-penetrant inhibitor, could reduce body weight in DIO mice, prompting us to pursue dedicated studies explicitly designed to confirm weight-loss effects under rigorously controlled conditions. To conclusively address whether CNS penetration is required for efficacy, we compared the centrally penetrant compound TN-783 with the peripherally restricted analog TN-101 (**Figure 3A**). Multiple doses of TN-783 were tested up to 50 mg/kg, while TN-101 was evaluated at 1 mg/kg (**Supplementary Table 3**). Lean animals were included at the 50 mg/kg dose of TN-783 to control for potential nonspecific effects of treatment at the highest effective dose selected for this design. In this experiment, animals were acclimated with twice-daily vehicle dosing for 14 days to habituate them to the regimen and allow body weight to stabilize. The mice were then randomized to treatment groups to ensure comparable baseline body weight and body composition (**Supplementary Figure 2A-B**).

**Figure 3.**
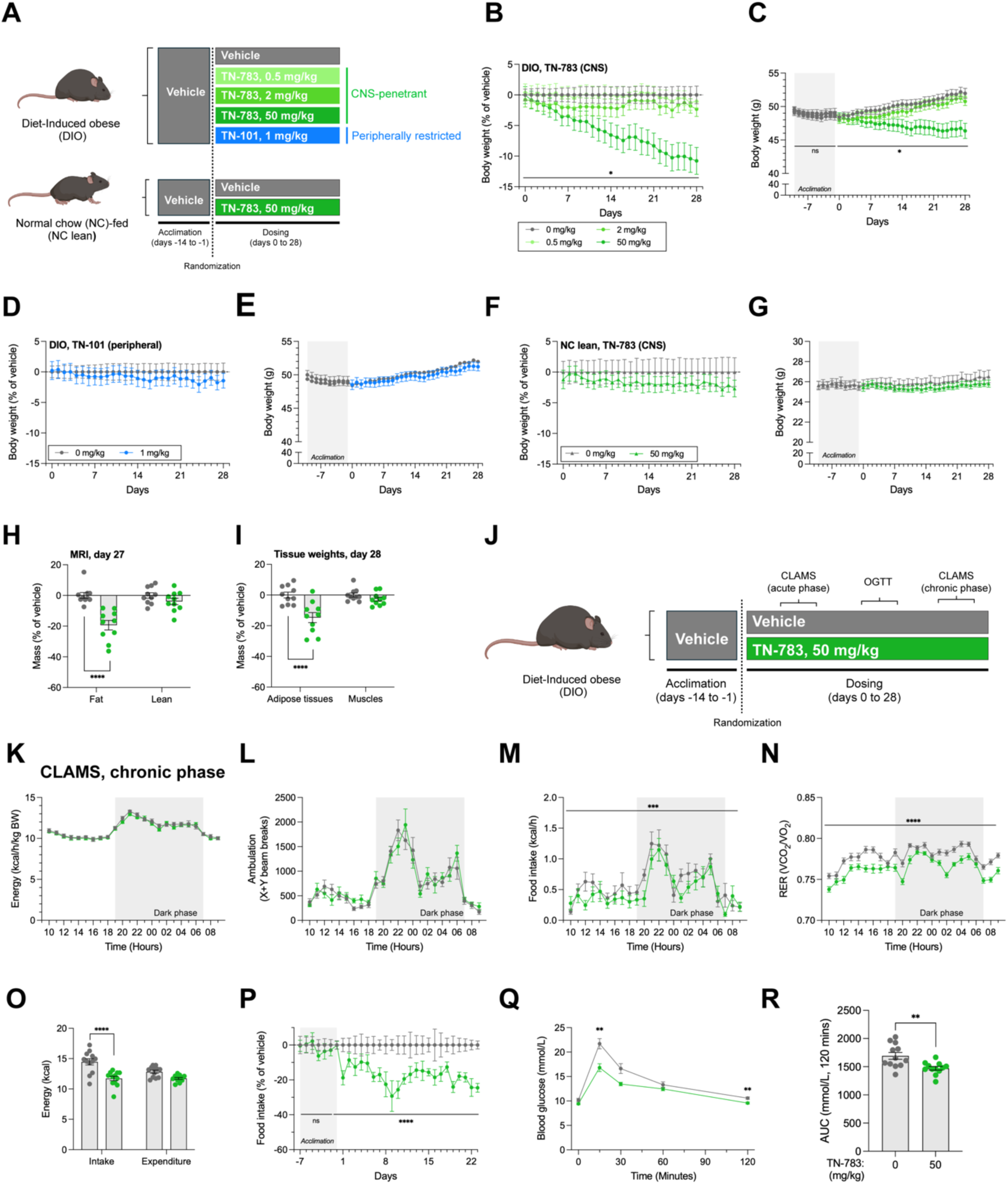
CNS-penetrant NLRP3 inhibitor induced weight loss in DIO mice. (**A**) Schematic of compound dosing in DIO and normal chow (NC)-fed lean mice. (**B-C**) Vehicle-adjusted and absolute body weight of DIO mice treated with TN-783. (**D-E**) Vehicle-adjusted and absolute body weight of DIO mice treated with TN-101. (**F-G**) Vehicle-adjusted and absolute body weight of lean mice treated with TN-783. Grey areas represent the acclimation period. (**H**) Fat and lean masses (vehicle-adjusted) after 27 days of TN-783 treatment in DIO mice assessed by MRI. (**I**) Masses of adipose tissues and muscles (vehicle-adjusted) after 28 days of TN-783 treatment in DIO mice. (**J**) Experimental schematic of TN-783 dosing study in DIO mice with Comprehensive Lab Animal Monitoring System (CLAMS) and oral glucose tolerance test (OGTT) measurements. (**K-N**) Energy expenditure (body weight-adjusted), physical activity, food intake, and respiratory exchange ratio (RER) over 24 hours in the chronic phase of TN-783 dosing. Grey areas indicate lights-off period. (**O**) Total energy intake and energy expenditure over the 24-hour period measured in the chronic phase of the study. (**P**) Daily food intake (vehicle-adjusted) over the acclimation and dosing periods. (**Q-R**) Blood glucose and area under the curve during the OGTT. Data are mean ± SEM of N=8-10/group (**A-I**) or N=12/group (**J–R**). Statistical significance for TN-783 effect was determined by a two-way ANOVA (**B-G, K-N, P**) followed by Šídák’s multiple comparison tests against vehicle-treated controls (**H-I**, **O**, **Q**) or Mann Whitney test (**R**) (*, p<0.05. **, p<0.01, ***, p<0.001, ****, p<0.0001).

Brain exposures were measured from terminal samples collected 1 hour after the final dose (**Supplementary Figure 2C**). At 50 mg/kg, TN-783 achieved unbound brain concentrations in both lean and DIO mice that exceeded the *in vitro* IC_90,u_ and aligned with levels predicted from plasma PK (**Supplementary Table 3**). At lower doses (0.5 and 2 mg/kg), unbound brain concentrations in DIO mice were near or below the IC_90,u_ level. In contrast, TN-101 administered at 1 mg/kg produced unbound brain levels below the limit of quantification (>16-fold below IC_90,u_), while exceeding IC_90_,_u_ coverage peripherally, confirming its restriction from the CNS.

We next examined whether these brain exposures translated into measurable effects on body weight in DIO and lean mice. In DIO animals, TN-783 produced a progressive and sustained reduction in body weight over 28 days at the 50 mg/kg dose, with no effect at lower doses (**Figure 3B-C**). Analyses performed both as percent of vehicle-normalized change and as absolute body weight yielded concordant results, supporting the robustness of the effect. By contrast, TN-101 had no impact on body weight in DIO mice (**Figure 3D-E**). Further, normal chow-fed lean mice treated with TN-783 at 50 mg/kg showed no weight change (**Figure 3F-G**). Collectively, our data established that CNS NLRP3 inhibition is required to drive weight loss. More specifically, this effect occurs only when central exposures exceed the IC_90,u_ and that this effect is specific to the obese state rather than a nonspecific consequence of the pharmacological agent.

To evaluate the impact of TN-783-induced weight loss on body composition, we performed MRI analysis after 27 days of dosing. Weight loss in TN-783-treated DIO animals was associated with a marked reduction in fat mass without significant reduction in lean mass (**Figure 3H, Supplementary Figure 2D**). This effect was confirmed by terminal tissue weights showing significantly decreased white adipose tissue depots, while representative muscle masses were preserved (**Figure 3I**, **Supplementary Figure 2E**). These effects of TN-783 were absent in NC lean animals treated at the same dose of 50 mg/kg (**Supplementary Figure 2G–H**), highlighting that TN-783 exerted its effect only in the context of obesity, with minimal impact under lean conditions. Food intake measurements showed that TN-783 induced a modest yet persistent reduction in caloric intake in DIO mice that was maintained throughout the dosing period and not observed in lean controls (**Supplementary Figure 2F, I**). Cumulative intake analysis (**Supplementary Figure 2J**) further showed that this sustained reduction brought food consumption level in DIO mice toward the range observed in NC lean animals.

The durability of the food intake suppression prompted us to next examine how TN-783 influences energy balance in more detail. To this end, we conducted comprehensive metabolic phenotyping using indirect calorimetry in DIO mice at both the acute and chronic phases of treatment on days 7-11 and 21-25, respectively (**Figure 3J; Supplementary Figure 3A**). Mice were acclimated to *b.i.d.* dosing for 14 days prior to study start, then assigned to treatment groups with matched baseline body weight and body composition (**Supplementary Figure 3B**). TN-783 was dosed twice daily at 50 mg/kg, the regimen previously shown to produce robust weight loss. In this study, TN-783 again induced progressive weight loss in DIO mice (**Supplementary Figure 3C-D**), and longitudinal MRI measurement confirmed the selective reduction in fat mass without significant alteration in lean mass (**Supplementary Figure 3E-G**).

To assess the relative contribution of energy expenditure and intake, animals were acclimated in Comprehensive Lab Animal Monitoring System (CLAMS) chambers for 48 hours prior to data collection, followed by a 48-hour recording window (24-hour averages shown). Energy expenditure per animal was modestly lowered in TN-783-treated mice at both the acute and chronic phases (**Supplementary Figure 3H, N)**. This decrease reflected the reduced body weight of TN-783-treated mice, as weight-normalized values were identical to those of controls (**Figure 3K, Supplementary Figures 3I**). Physical activity was similarly unchanged by TN-783 treatment (**Figure 3L**; **Supplementary Figure 3J**). In contrast, food intake was consistently reduced in TN-783-treated DIO mice across both the acute and chronic phases (**Figure 3M**; **Supplementary Figure 3K**). This suppression was accompanied by a sustained lowering of respiratory exchange ratio (RER; **Figure 3N**; **Supplementary Figure 3L**), a shift often observed when food intake is reduced and reflecting greater reliance on lipid utilization (32). Analysis of total energy balance showed that TN-783 treatment reduced energy intake without altering energy expenditure (**Figure 3O**, **Supplementary Figure 3M**), shifting these DIO animals from a net positive balance under vehicle to approximate equilibrium. As a result, TN-783-treated mice maintained a lower energy balance than controls during both the acute and chronic phases of treatment. Longitudinal monitoring of food intake across the full study period (**Figure 3P**; **Supplementary Figure 3O**) further confirmed this effect: TN-783 reduced food intake immediately upon initiation of dosing, and the reduction was sustained throughout the 28-day study.

An oral glucose tolerance test (OGTT) was carried out on day 15 to determine whether TN-783 treatment improved metabolic control. TN-783-treated DIO mice displayed lower blood glucose excursions compared to vehicle controls following the glucose challenge (**Figure 3Q**). This improvement was further reflected by a significant reduction in the area under the curve (AUC; **Figure 3R**), indicating that enhanced glucose tolerance emerged in parallel with the ongoing loss of body weight and fat mass.

Together, these studies demonstrated that TN-783 induced fat-selective weight loss in obese mice by persistently suppressing food intake without affecting energy expenditure or activity. The resulting improvements in body composition were accompanied by enhanced glucose tolerance, further underscoring the therapeutic potential of CNS-penetrant NLRP3 inhibitors in obesity and its comorbidities.

### TN-783 attenuated hypothalamic gliosis and partially restored obesity associated molecular dysregulation

Because the weight-lowering effects of TN-783 required CNS penetration and were mediated through suppression of feeding, we focused our mechanistic analysis on the hypothalamus, a key brain region that controls energy homeostasis. The hypothalamus integrates hormonal and neuronal cues to regulate appetite and metabolism (33) and gliosis within this region has previously been reported in obesity across both animal models and humans (16, 20). These observations led us to hypothesize that TN-783 may alleviate hypothalamic neuroinflammation.

Precise delineation of hypothalamic nuclei is technically demanding and can be challenging even for experienced investigators. To manage this complexity, we employed AgRP immunoreactivity as a structural marker: AgRP is expressed in orexigenic neurons whose cell bodies reside in the hypothalamic arcuate nucleus (ARC). Within the hypothalamus, AgRP neurons extend projections within the ARC and to the dorsomedial hypothalamus (DMH), but are absent from the ventromedial hypothalamus (VMH), thereby serving as a robust supplementary marker that complements conventional neuroanatomical landmarks (**Supplementary Figure 4**). To assess glial activation, we measured Iba1, a marker for microglia, and GFAP, a marker for reactive astrocytes, since increases in these markers are indicative of neuroinflammation. GFAP immunoreactivity in the ARC was not significantly different between groups (**Supplementary Figure 5**). In contrast, DIO mice exhibited a significant increase in Iba1 immunoreactivity in the ARC, consistent with microglial activation, while levels in the VMH and DMH were not different (**Figure 4A-B**). Importantly, treatment of DIO mice with TN-783 significantly reduced Iba1 immunoreactivity in the ARC, restoring the values toward those observed in NC lean animals. Collectively, these results demonstrated that DIO led to significant hypothalamic microgliosis, and that TN-783 selectively attenuated this response in the ARC, a critical center for appetite and energy balance regulation.

**Figure 4.**
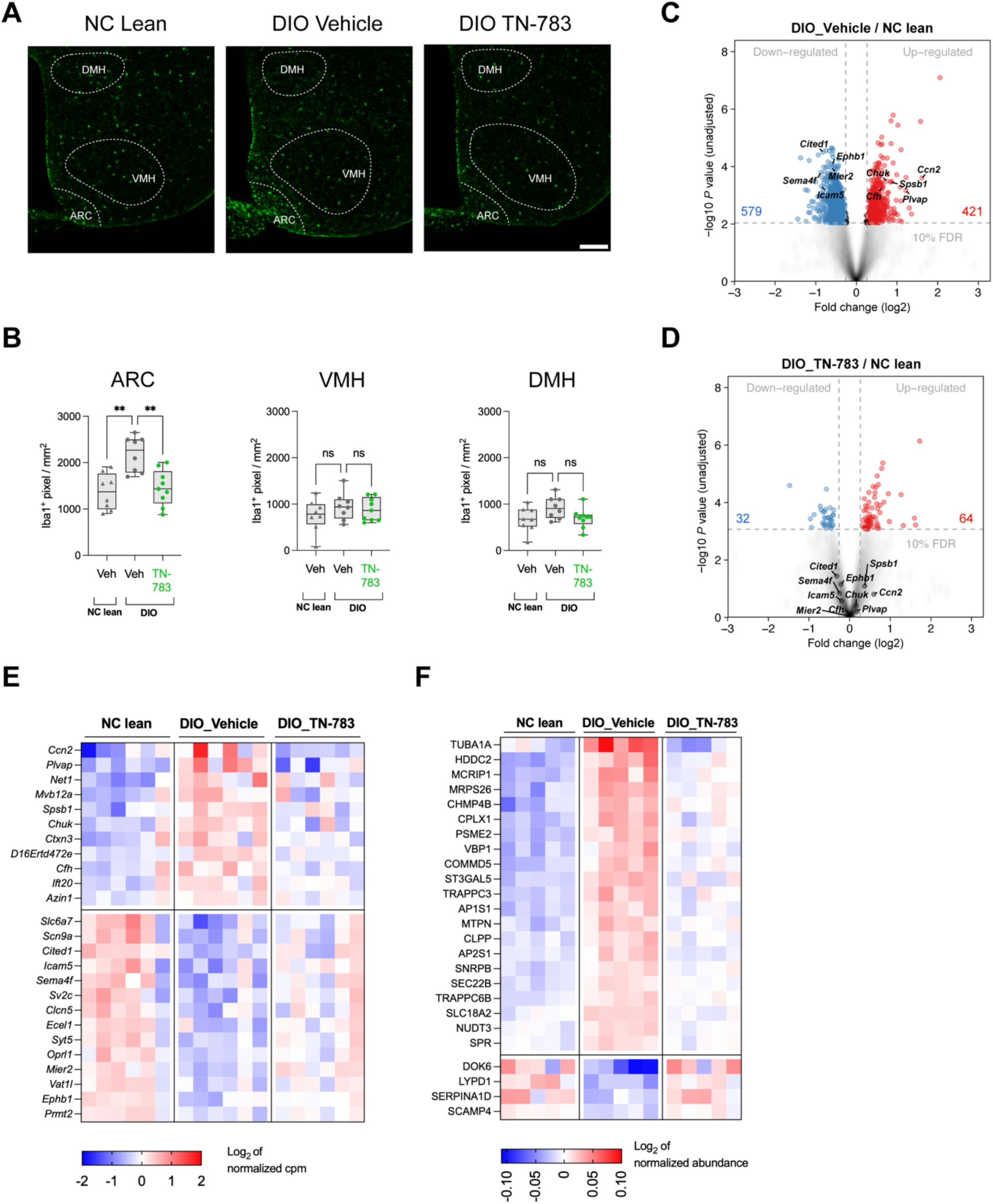
TN-783 reduced microgliosis marker Iba1 in the hypothalamic arcuate nucleus (ARC) in association with partial rescue of DIO-induced transcriptomic and proteomic changes. (**A**) Representative sections showing Iba1 immunoreactivity in the hypothalamus of vehicle-treated NC lean control, vehicle- and TN-783-treated DIO animals. Scale bar: 100 µm. ARC: arcuate nucleus, VMH: ventromedial hypothalamus, DMH: dorsomedial hypothalamus. Scale bar: 100 µm (**B**) Quantification of Iba1 immunostaining. Data are shown as median with interquartile range (IQR); whiskers indicate min–max. Statistical significance for TN-783 effect was determined by a one-way ANOVA followed by Dunnett’s multiple comparison tests against vehicle-treated DIO controls (*, p<0.05. **, p<0.01). (**C-D**) Volcano plots showing transcriptomic changes in the hypothalamus of DIO mice relative to NC lean controls (**C**) and TN-783-treated DIO mice relative to NC lean controls (**D**). Genes with false discovery rate (FDR, adjusted p value) < 10% and absolute fold change > 20% are marked by red (upregulated) and blue (downregulated) circles. Examples of genes significantly altered in DIO relative to lean but rendered insignificant with TN-783 treatment are labeled. (**E-F**) Heatmaps for the top 25 hits (see **Supplementary Figure 6C** & **6E** for selection criteria) demonstrating alterations in DIO relative to NC lean controls with rescue by TN-783 treatment from bulk RNA sequencing (**E**) and proteomics (**F**) of hypothalamus.

Previous studies also reported profound alterations at both transcriptional and protein levels in the hypothalamus of mice fed a high-fat diet (34–38). To determine whether TN-783 can ameliorate DIO-associated hypothalamic dysregulation besides gliosis, we performed bulk RNA sequencing on hypothalami dissected from NC lean mice and DIO mice treated with vehicle (DIO_Vehicle) or TN-783 (DIO_TN-783) for 4 weeks. All samples exhibited clear expression of hypothalamic marker genes (39), with minimal expression of a cerebellum-specific control markers (40) (**Supplementary Figure 6A**). Consistent with previous reports, DIO_Vehicle mice displayed an extensive set of differentially expressed genes (DEGs), including 421 significantly upregulated and 579 significantly downregulated (FDR < 10%, absolute fold change > 20%) (**Figure 4C**). The key pathways perturbed by DIO were enriched for cellular metabolism, inflammation, and signaling (**Supplementary Table 4**). Notably, treatment with TN-783 reversed the expression patterns of many DIO-induced DEGs (**Supplementary Figure 6B,** compared with **Figure 4C**), reducing the total number of DEGs to 96 (64 significantly upregulated, 32 significantly downregulated) (**Figure 4D**). These findings suggested that TN-783 markedly attenuated transcriptional dysregulation in the hypothalamus of DIO mice. A heatmap of the top 25 DEGs rescued by TN-783 (see **Supplementary Figure 6C** for selection algorithm) showed that genes upregulated or downregulated in DIO_Vehicle mice were restored toward the expression profile of NC lean controls (**Figure 4E**). Among the top genes rescued by TN-783 were those involved in immune functions (e.g., *Cfh* [complement factor H, regulator of the complement cascade], *Chuk* [IKKα, regulator of NF-κB signaling]), neuronal signaling (e.g., *Sema4f*, *Icam5*), and transcriptional regulation (e.g., *Mier2*, *Cited1*) (**Supplementary Figure 6D**).

We next performed proteomics on hypothalami isolated from a 14-day study with the same three treatment groups to assess the impact of obesity and TN-783 treatment at the protein level. DIO mice demonstrated significant changes in the relative abundance of >100 proteins (103 significantly up and 19 significantly down at FDR < 10% and absolute fold change > 20%) (**Supplementary Figure 6E**). Similar to its effect on the transcriptome, TN-783 treatment attenuated many of these changes, with the top 25 proteins rescued by TN-783 illustrated in the heatmap (**Figure 4F, Supplementary Figure 6F**) and individual plots of the top 8 proteins shown in **Supplementary Figure 6G**. These proteins are functionally enriched in trafficking (e.g., TUBA1A, TRAPPC3, TRAPPC6B, AP2S1, AP1S1, CHMP4B), neurotransmission and neuronal signaling (e.g., CPLX, SEC22B, LYPD1, DOK6), and immune regulation (e.g., PSME2, SERPINA1D, COMMD5). Taken together, our results demonstrated that DIO promoted pathological shifts in the hypothalamic microenvironment, characterized by microglial activation and broad molecular dysregulation.

### CNS-penetrant NLRP3 inhibitor TN-783 enhanced semaglutide efficacy, restarted weight loss after plateau, and maintained weight reduction following withdrawal

Because GLP-1RAs represent the current standard of care in obesity, we investigated whether central NLRP3 inhibition could enhance or extend their therapeutic benefit. We therefore examined the ability of TN-783 to augment semaglutide-induced weight loss and to preserve weight reduction after semaglutide withdrawal (**Figure 5A-I**). Mice were first acclimated by twice-daily oral gavage with the vehicle solution for TN-783 and once-daily subcutaneous injection with the vehicle for semaglutide for 13 consecutive days. After this mock-dosing period, animals were randomized into treatment groups to ensure comparable baseline body weight and body composition, as in previous studies. The study then proceeded through weight loss induction, cross over, and maintenance phases. Semaglutide was administered at 0.0041 mg/kg (1 nmol/kg), a dose known to induce robust, but submaximal weight loss (41), thereby allowing us to test whether TN-783 could further improve efficacy.

**Figure 5.**
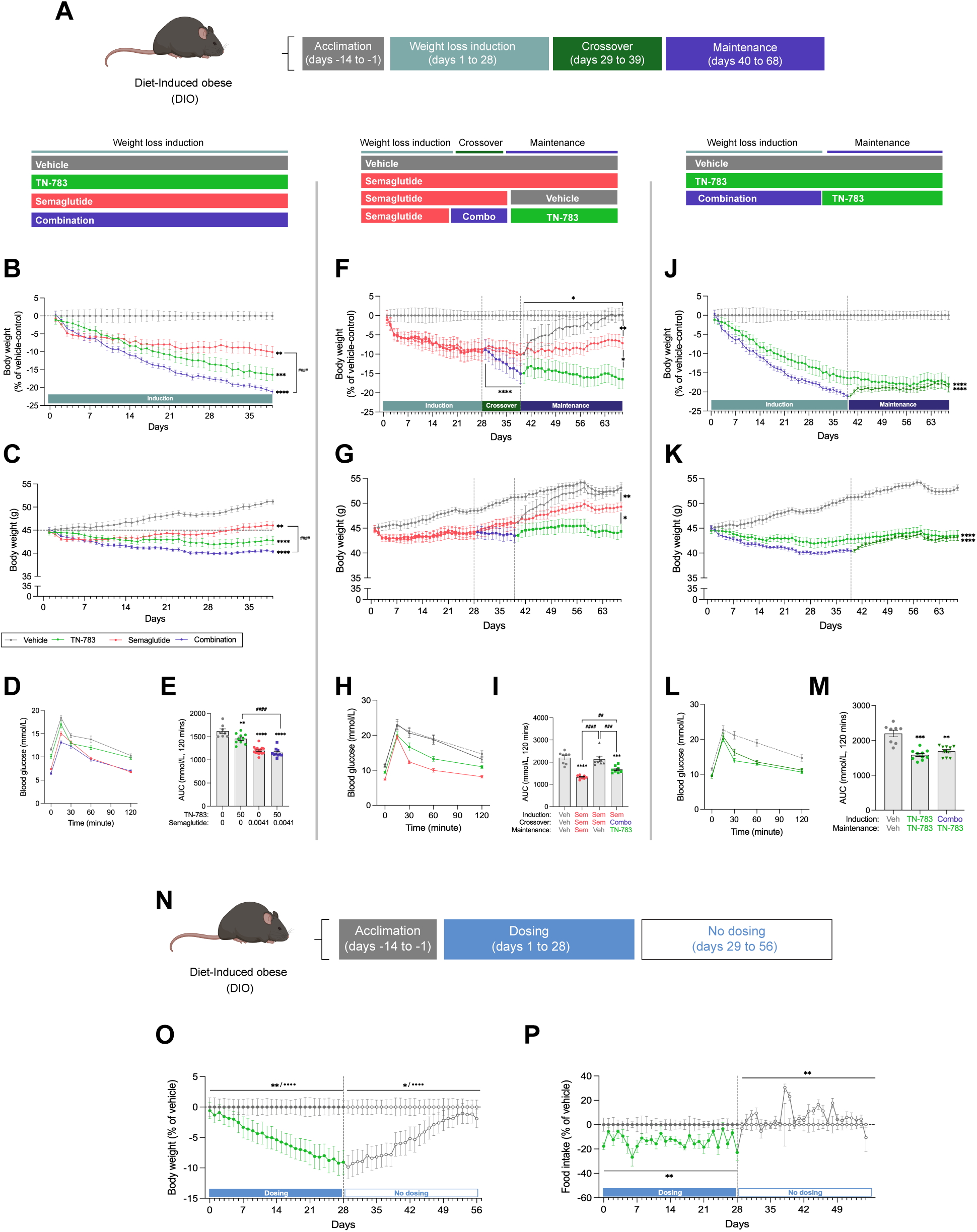
TN-783 potentiated semaglutide-induced weight loss, enabled durable maintenance after semaglutide withdrawal, and showed reversibility consistent with an on-target effect. (**A**) Study schematic of TN-783 and semaglutide dosing in DIO mice. Following acclimation, mice underwent a weight loss induction phase (days 1–28), a crossover/extension period (days 29–39), and a maintenance phase (days 40–66). (**B-E**) Body weight (vehicle-adjusted and absolute) and OGTT following treatment with vehicle, TN-783, semaglutide, or their combination during the weight loss induction period. (**F-I**) Body weight (vehicle-adjusted and absolute) and OGTT following treatment with vehicle, semaglutide, semaglutide followed by vehicle, or semaglutide followed by combination treatment with TN-783 in the crossover phase and TN-783 monotherapy during the maintenance phase. (**J-M**) Body weight (vehicle-adjusted and absolute) and OGTT following treatment with vehicle, TN-783 monotherapy, or its combination with semaglutide during the weight loss induction phase followed by TN-783 monotherapy during the maintenance period. (**N**) Study schematic of TN-783 dosing in DIO mice during weight loss induction (days 1–28) followed by withdrawal (days 29–56). (**O-P**) Body weight (vehicle-adjusted) and food intake following treatment with TN-783 during the weight loss induction period and after treatment withdrawal. Data are mean ± SEM of N=8-16/group (**B-E**), N=8-10/group (**F-M**) or N=7-8/group (**N-P**). Statistical significance for TN-783 effect was determined by a two-way ANOVA (**B-C, J-K, O-P**) followed by multiple comparison tests as indicated (**F-G**) or one-way ANOVA followed by Tukey’s multiple comparison tests against vehicle-treated DIO controls or as indicated (**E, I, M**). Asterisks (*) indicate a significant main effect of drug, whereas bullet point (•) indicate a significant drug x time interaction (**/••, p<0.01, ***/•••, p<0.001, ****/••••, p<0.0001).

Consistent with published data, semaglutide treatment produced a rapid and robust reduction in body weight relative to vehicle controls, which then plateaued after approximately two weeks (**Figure 5B-C**). In contrast, TN-783 monotherapy drove a more gradual yet progressive weight loss that continued throughout the treatment period, reproducing the effect observed in prior studies (**Figures 3B-C** & **Supplementary Figures 3D-D**). Importantly, combining TN-783 with semaglutide not only resulted in a greater magnitude of weight loss than either monotherapy, but also a sustained and progressive trajectory, illustrating that NLRP3 inhibition can enhance and extend GLP-1-based efficacy. Analysis of individual weight loss curves revealed substantial variability in response to either monotherapy (**Supplementary Figure 7A-G**), whereas the combination therapy produced a more uniform weight-loss profile across animals (**Supplementary Figure 7H-I**). In addition, mice receiving the combination therapy showed improved glucose tolerance relative to vehicle and TN-783 alone, with significantly lower glucose excursions during an oral glucose tolerance test and reduced area under the curve (**Figure 5D-E**).

While semaglutide induces substantial weight loss, its clinical utility is limited by a plateau in efficacy and by tolerability issues that impede long-term adherence. We therefore asked if TN-783 could reinitiate weight loss once the semaglutide response had plateaued and sustain weight reduction following semaglutide discontinuation. As shown in the design schematic above **Figure 5F**, animals were first treated with semaglutide, then randomized to different maintenance regimens. Addition of TN-783 during the crossover phase, after semaglutide-induced weight loss had plateaued, rapidly triggered further reductions in body weight (**Figure 5F-G**), mirroring the additive effect observed in the induction phase with combination therapy (**Figure 5B-C**). This renewed weight loss indicated that NLRP3 inhibition could overcome resistance to semaglutide and reinstate its weight-loss effect after the initial response had diminished. Transition of this group to TN-783 monotherapy preserved the reduced body weight with minimal rebound. In contrast, mice switched to vehicle rapidly regained weight, and those maintained on semaglutide showed a stable profile without further reduction. By the end of the study, the group treated with TN-783 in the maintenance phase reached a lower body weight than either comparator group (**Figure 5F-G**). These data demonstrated that TN-783 alone can sustain and maintain weight loss following semaglutide withdrawal.

To assess whether these maintenance effects extended to metabolic function, we performed an OGTT on day 58. Mice that continued semaglutide treatment showed improved glucose tolerance compared to vehicle controls (**Figure 5H-I**), whereas animals switched from semaglutide to vehicle exhibited glycemic excursions indistinguishable from vehicle-only controls, suggesting a complete loss of benefit. By contrast, mice transitioned to TN-783 monotherapy displayed reduced glucose excursions and lower AUC relative to those maintained on vehicle throughout and those switched from semaglutide to vehicle, indicating partial preservation of glycemic control by central NLRP3 inhibition despite semaglutide withdrawal.

The durability of TN-783-induced weight loss was further assessed in animals treated with either continuous monotherapy or a regimen involving combination therapy during induction followed by TN-783 alone during maintenance (**Figure 5J-K**). In both groups, body weight declined significantly during the induction period and was stably maintained throughout the maintenance phase under TN-783 monotherapy. By study end, the two regimens yielded comparable body weights. Mice treated with either continuous TN-783 or the combination-to-monotherapy sequence displayed improved glucose tolerance compared with vehicle controls, as reflected by lower glycemic excursions during the challenge and significantly reduced AUC values (**Figure 5L-M).** No significant differences were observed between the two TN-783 regimens. These findings demonstrated that our CNS-penetrant NLRP3 inhibitor, whether administered as continuous monotherapy or following initial combination with semaglutide, supports durable weight loss and sustained metabolic improvement in DIO mice.

Finally, we assessed whether the effects of TN-783 were reversible upon treatment cessation (**Figure 5N**, **Supplementary Figure 8A**). After 28 days of TN-783 dosing, animals displayed robust reduction in body weight relative to vehicle controls as expected (**Figure 5O**, **Supplementary Figure 8B**). Withdrawal of TN-783 led to a rapid rebound in food intake along with progressive weight regain, ultimately converging toward the trajectory of vehicle-treated mice (**Figure 5O-P**, **Supplementary Figure 8B-C**). These results indicated that the weight-loss effect of TN-783 is fully reversible and dependent on continued target engagement. Collectively, these results demonstrated that central NLRP3 inhibition offers a novel, safe, and versatile approach to obesity treatment, with potential efficacy as a monotherapy and the capacity to both extend and enhance the benefits of existing therapies.

## DISCUSSION

Our study identified central NLRP3 inhibition as a mechanistically distinct therapeutic approach from incretin-based therapies for obesity. We demonstrated that efficacy required direct CNS engagement, with efficacy achieved only when unbound brain concentrations exceeded the *in vitro* IC_90,u_ threshold for target inhibition. TN-783 induced progressive, fat-selective weight loss through sustained suppression of food intake without altering energy expenditure. At the mechanistic level, TN-783 corrected obesity-associated microgliosis in the hypothalamic ARC. Associated with this, transcriptomic and proteomic profiling of the hypothalamus revealed partial reversal of obesity-associated molecular changes, particularly in pathways related to inflammation and neuronal signaling. These findings provided mechanistic insight into why CNS target engagement is required for NLRP3 inhibition to confer metabolic benefit. In the therapeutic context, TN-783 enhanced the effects of semaglutide when co-administered and, critically, TN-783 reinitiated weight loss after semaglutide’s effect had plateaued, demonstrating central NLRP3 inhibition as an effective add-on to GLP-1RA therapy. Moreover, TN-783 preserved weight loss after semaglutide withdrawal and demonstrated full reversibility upon discontinuation. Together, these findings define the mechanistic basis and highlight the therapeutic potential of a CNS-penetrant NLRP3 inhibitor in obesity.

The hypothalamus has long been recognized as a central regulator of energy homeostasis. This region is particularly vulnerable to SFA, such as palmitate, which rapidly accumulate in the mediobasal hypothalamus following high-SFA intake (42). Supporting a causal link between dietary SFA and NLRP3 activation, palmitate exposure triggered robust secretion of NLRP3-dependent cytokines IL-1β and IL-18 from macrophage-like cells and this inflammatory response was potently suppressed by NLRP3 inhibitors. These cellular findings provide a mechanistic link between dietary lipids and tissue-level consequences of chronic high-fat feeding. Inflammation of the mediobasal hypothalamus in DIO is well documented in rodents and has also been observed in humans by MRI and histological analyses (43–46). Consistent with prior studies (42, 46–48) we found that DIO was associated with selective microgliosis in the ARC, a key appetite-regulating region in the mediobasal hypothalamus, and that treatment with the CNS-penetrant NLRP3 inhibitor TN-783 rescued this gliosis, restoring microglial reactivity toward levels observed in lean animals. Besides microgliosis, DIO also induces widespread transcriptomic and proteomic alterations in the hypothalamus (49–53). Consistent with these prior findings, our transcriptomic and proteomic analyses of the DIO hypothalamus revealed robust changes in inflammatory and metabolic pathways, as well as dysregulation of neuronal signaling and trafficking proteins, demonstrating the breadth of molecular disruptions in the hypothalamus during obesity. Remarkably, TN-783 dampened many DIO-induced DEGs and protein-level changes, supporting a mechanistic role of NLRP3 in hypothalamic dysregulation. The attenuation of changes in inflammatory genes and proteins underscores the central role of the NLRP3 inflammasome in neuroinflammation, while the rescue of neuronal signaling and neurotransmission pathways likely reflects secondary benefits of reduced microgliosis. This interpretation aligns with reports showing that resolving hypothalamic inflammation, whether by depleting microglia, genetic silencing of inflammatory signaling (IKKβ/NF-κB, JNK pathways), or direct anti-inflammatory blockade (inhibition of TLR4/TNFα) can alleviate neuronal stress (18, 19, 45, 54, 55). Together, these findings highlight the central role of NLRP3 in hypothalamic neuroinflammation and demonstrate that central NLRP3 inhibition reverses both cellular and molecular hallmarks of obesity-associated hypothalamic dysfunction.

To understand why resolving neuroinflammation leads to reduced food intake, we focused on the physiological function of the ARC, which contains first-order neurons that sense peripheral and central signals of energy status and orchestrate feeding behavior through interconnected circuits (56–58). Neuroinflammation within this nucleus has been proposed to compromise the functional integrity of these neurons, impairing their ability to respond to regulatory cues (42, 47–48). DIO in both mouse models and humans is well known to be associated with resistance to leptin, an adipocyte-derived hormone. Leptin normally contributes to homeostatic regulation of energy balance by inhibiting orexigenic neurons and stimulating anorexigenic neurons in the ARC (59). Prior work showed that targeting neuroinflammation by ablating microglia in DIO mice restored leptin sensitivity, resulting in reduced food intake and protection against further weight gain (19). This work provided direct evidence that microglia activity can impair satiety signaling and thereby promote obesity. More broadly, microglia-driven disruption of neuronal function is a common feature across disease contexts, including Alzheimer’s disease (60, 61), multiple sclerosis (62), and traumatic brain injury (63), where microglial activation has been implicated in synaptic remodeling, neurotransmission deficits, and altered circuit connectivity leading to functional impairment (62, 64, 65). These observations highlight how inflammatory remodeling of neuronal circuits can compromise network fidelity across contexts, raising the question of how local microenvironmental features of the ARC makes it especially vulnerable in obesity. The ARC lies adjacent to the median eminence, a circumventricular organ that lacks a classical BBB. This arrangement allows circulating hormones, nutrients, and metabolites to directly access subsets of ARC neurons, while other neuronal populations within the same nucleus remain BBB-protected (58, 66). Such mixed exposure creates a heterogeneous environment in which some neuronal subtypes are directly exposed to circulating dietary and metabolic signals, while others rely on indirect or glial-mediated sensing. In the context of chronic high-fat feeding, this dual architecture may render the ARC particularly susceptible to inflammatory remodeling: neurons at the BBB-free interface are continuously exposed to elevated lipids and cytokines, while surrounding microglia integrate both central and peripheral stress signals. This unique positioning of the ARC raises an important pharmacologic question: if subsets of ARC neurons are directly exposed to circulating factors, why is a CNS-penetrant NLRP3 inhibitor required to achieve metabolic benefit? We propose that the signaling integrity of neurocircuitry within BBB-protected networks of the ARC is critical for the coordinated regulation of energy homeostasis and feeding behavior. Within this nucleus, first-order orexigenic AgRP/NPY and anorexigenic proopiomelanocortin (POMC) neurons form the fundamental circuit governing feeding behavior (57, 67–69). Notably, nonspecific lesioning of the ARC region located outside the BBB in adult mice produces little or no effect on food intake or body weight, indicating that effective regulation of feeding behavior depends critically on the integrity of neurons and circuits situated behind the barrier (58). Another possibility is that other brain regions shielded by the BBB may also undergo inflammatory remodeling in obesity such that broader CNS engagement is required to fully restore homeostatic function. These considerations suggest that the metabolic benefits of NLRP3 inhibition ultimately reflect the ability of a highly CNS-penetrant inhibitor to recalibrate hypothalamic circuits more broadly, thereby restoring the neuronal fidelity required for long term regulation of energy balance.

Beyond its efficacy as a monotherapy, we observed that central NLRP3 inhibition not only enhanced the effect of semaglutide but also reinitiated weight loss after semaglutide weight loss had plateaued. GLP-1RAs are known to drive rapid appetite suppression but typically plateau with continued treatment (2, 70), suggesting the emergence of resistance. One possible explanation is that persistent and unresolved hypothalamic inflammation impairs the ability of neurons in the ARC to sense and integrate satiety signals, both endogenous (such as leptin) and exogenous (such as GLP-1 analogs), thereby acting as a barrier to limit continued incretin response. Under this framework, incretin therapy amplifies satiety inputs, but the inflamed hypothalamus has diminished capacity to respond to them, leading to a plateau in weight loss despite persistent excess adiposity. While GLP-1 receptor agonists, including semaglutide, have been reported to exert anti-inflammatory effects (71–74), our findings suggest that hypothalamic neuroinflammation nonetheless persists during semaglutide treatment and remains responsive to NLRP3 inhibition for resolution. By suppressing this neuroinflammation, NLRP3 inhibition appeared to restore neuronal responsiveness, effectively lowering the resistance to satiety hormones and allowing feeding circuits to remain sensitive to ongoing signals. This mechanistic framework suggests that the benefit of NLRP3 inhibition may extend beyond GLP-1 to other satiety hormone pathways, including glucagon, GIP, amylin, and PYY, by improving the brain’s ability to sense and respond to these cues. Together, these complimentary mechanisms highlight the potential of NLRP3 inhibition to extend and stabilize the therapeutic benefits of incretins in clinical practice.

Taken together, our findings provide important insights into the specificity, reversibility, and translational potential of CNS-penetrant NLRP3 inhibitor in obesity. TN-783 produced weight loss in obese, but not lean mice, and this was achieved while preserving lean muscle mass. This selective profile addresses a long-standing challenge for obesity pharmacotherapy where loss of lean tissue often accompanies fat reduction. The profile of TN-783 was also distinct from semaglutide in that weight loss was progressive rather than abrupt, and was driven by modest, sustained suppression of food intake, a dynamic typically associated with healthier body composition and less drastic muscle loss than therapies that induce rapid catabolism (75). Furthermore, withdrawal of TN-783 led to rapid weight regain, consistent with the notion that ongoing dietary lipid stress continually activates hypothalamic microglia and that sustained target CNS engagement is required to maintain benefit. Taken together, these findings position a CNS-penetrant NLRP3 inhibitor as a promising therapeutic strategy that restores hypothalamic function, achieves durable and fat-selective weight loss, and complements existing treatments. More broadly, these results underscore the importance of targeting neuroimmune pathways in obesity, and provide a framework for clinical translation, where CNS NLRP3 inhibition could be deployed as both a monotherapy and in combination with established agents.

## ACKNOWLEDGEMENT

We thank Cathal S. Mahon for coordinating the Cryo-EM structure of TN-551 with NLRP3/NEK7, Ahmed El Sherbeni for technical support, Kate Stuart for insightful discussion on the manuscript, and Gilbert Di Paolo for his valuable scientific insights on the project.

## MATERIALS AND METHODS

### Compound synthesis and characterization

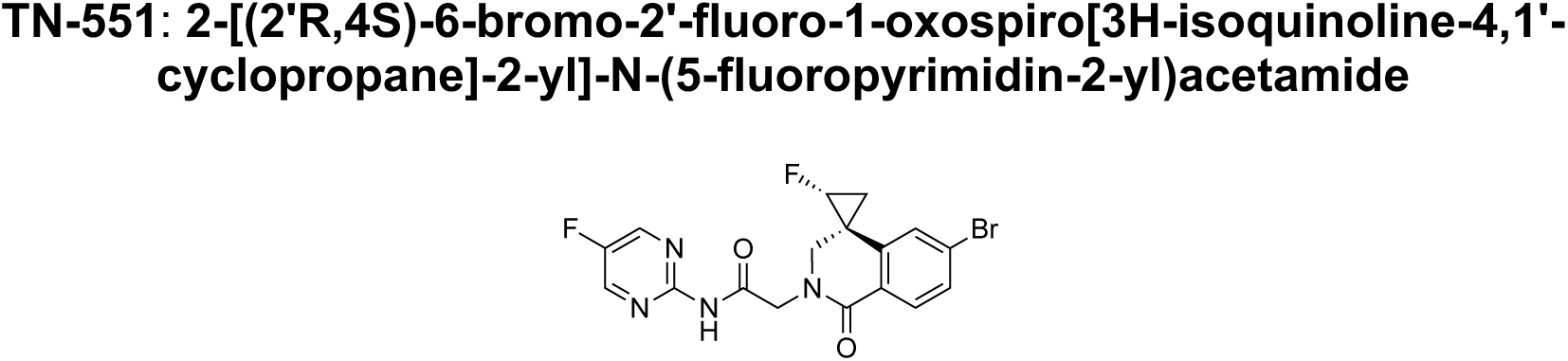

TN-551 can be prepared according to the procedures outlined in WO2022/232632.

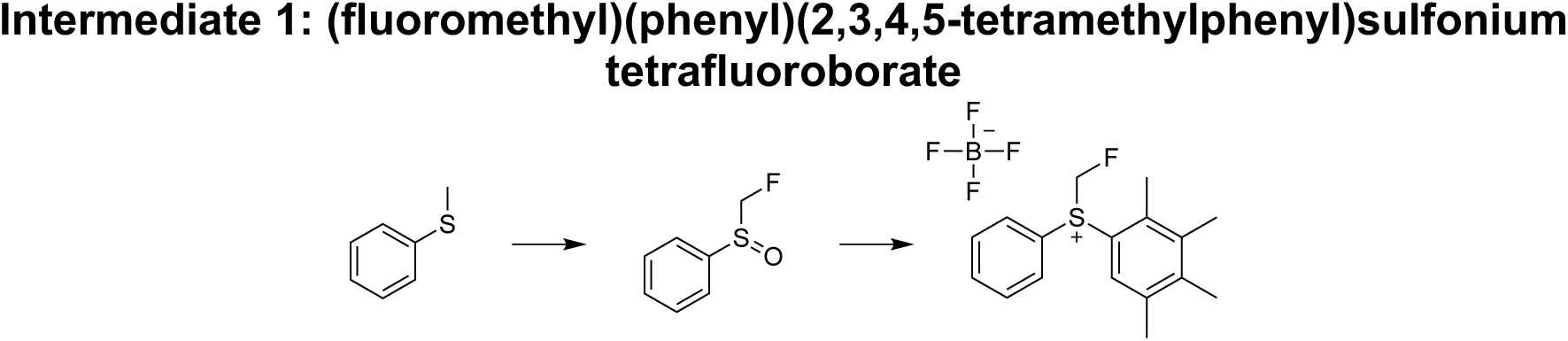

#### (fluoromethyl)sulfinyl)benzene

To a suspension of SelectFluor (463.5 g, 1.31 mol) in MeCN (1.5 L) at 0 °C was added a solution of methyl(phenyl)sulfane (130.0 g, 1.05 mol) in MeCN (150 mL) over 10 min. Et_3_N (132.4 g, 1.31 mol) was then added to the mixture at 0 °C. The reaction mixture was allowed to warm to 20 °C and stirred for 16 h. Two batches of the above were run in parallel and combined for workup. The combined batches were diluted with water (1000 mL) and extracted with DCM (3 × 500 mL). The combined organic layers were dried over anhydrous Na_2_SO_4_, filtered, and concentrated under reduced pressure. The crude residue was dissolved in MeOH (2000 mL), diluted with water (200 mL), and cooled to 0 °C. NBS (372.6 g, 2.09 mol) was carefully added, and the mixture was stirred for 16 h at 15 °C. The reaction mixture was diluted with 10% Na_2_SO_3_ solution (200 mL) followed by sat. aq. NaHCO_3_ until pH = 7. Methanol was removed under reduced pressure, and the remaining aqueous phase was extracted with DCM (3 × 500 mL). The combined organic layers were washed with brine (500 mL), dried over anhydrous Na_2_SO_4_, filtered, and concentrated under reduced pressure.

#### (fluoromethyl)(phenyl)(2,3,4,5-tetramethylphenyl)sulfonium tetrafluoroborate (Intermediate 1)

To a solution of (fluoromethyl)sulfinyl)benzene (10.0 g, 63.2 mmol) in diisopropyl ether (100 mL) at -10 °C were added 1,2,3,4-tetramethylbenzene (7.64 g, 56.9 mmol) and Tf_2_O (17.8 g, 63.2 mmol). The reaction mixture was stirred at 20 °C for 1 h. The reaction mixture was filtered, and the resulting solid was dissolved in DCM (200 mL) and washed with aq. NaBF_4_ (1 M, 6 × 200 mL). The organic phase was dried over anhydrous Na_2_SO_4_, filtered, and concentrated under reduced pressure. The residue was triturated with MTBE at 20 °C for 30 min and then filtered to provide the desired product. LCMS: *m/z* = 275.2 [M+H]^+^.

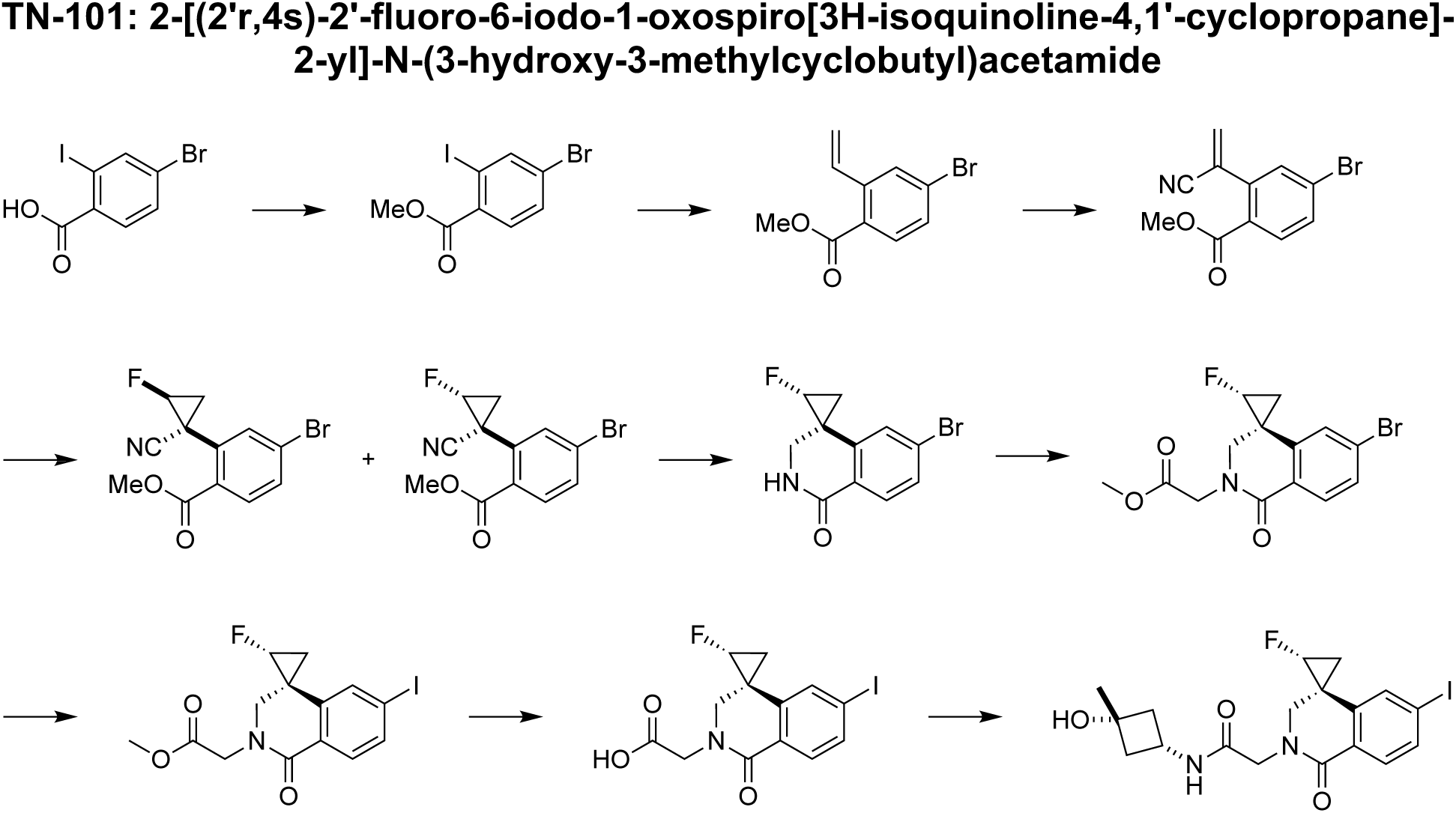

#### Methyl 4-bromo-2-iodobenzoate

To a solution of 4-bromo-2-iodo-benzoic acid (10.0 g, 30.6 mmol) in MeOH (100 mL) at 0 °C was added conc. H_2_SO_4_ (15 g, 153 mmol). The reaction mixture was stirred at 80 °C for 5 h. The reaction mixture was poured into water (300 mL) and extracted with EtOAc (3 × 100 mL). The combined organic layers were washed with sat. aq. NaHCO_3_ (3 × 50 mL), dried over anhydrous Na_2_SO_4_, filtered, and concentrated under reduced pressure to provide a crude residue that was used directly. ^1^H NMR (400 MHz, CDCl_3_): δ 8.18 (s, 1H), 7.70 (d, *J* = 8.4 Hz, 1H), 7.55 (dd, *J* = 0.8, 8.0 Hz, 1H), 3.93 (s, 3H).

#### Methyl 4-bromo-2-vinylbenzoate

To a mixture of methyl 4-bromo-2-iodo-benzoate (6.0 g, 17.6 mmol), potassium vinyltrifluoroborate (2.4 g, 17.6 mmol), and CsF (8.0 g, 52.8 mmol) in 1,4-dioxane (100 mL) was added Pd(dppf)Cl_2_ (1.3 g, 1.76 mmol). The reaction mixture was stirred at 90 °C for 16 h. The reaction mixture was concentrated under reduced pressure. The residue was purified by silica gel column chromatography. ^1^H NMR (400 MHz, CDCl_3_): δ 7.77 (d, *J* = 8.4 Hz, 1H), 7.72 (d, *J* = 2.0 Hz, 1H), 7.49-7.37 (m, 2H), 5.66 (d, *J* = 17.2 Hz, 1H), 5.41 (d, *J* = 10.8 Hz, 1H), 3.90 (s, 3H).

#### Methyl 4-bromo-2-(1-cyanovinyl)benzoate

To a mixture of Cu_2_O (345 mg, 2.41 mmol), 6,6’-dimethyl-2,2’-bipyridine (444 mg, 2.41 mmol), and Selectfluor (6.4 g, 18 mmol) in acetone (20 mL) and water (10 mL) were added methyl 4-bromo-2-vinylbenzoate (2.9 g, 12 mmol) and TMSCN (2.4 g, 24.1 mmol). The reaction mixture was stirred at 20 °C for 16 h. The reaction mixture was poured into aq. NaHCO_3_ (1 M, 30 mL) and extracted with EtOAc (3 × 30 mL). The combined organic layers were washed with brine (30 mL), dried over anhydrous Na_2_SO_4_, filtered, and concentrated under reduced pressure. The residue was purified by silica gel column chromatography. ^1^H NMR (400 MHz, CDCl_3_): δ 7.89 (d, *J* = 8.4 Hz, 1H), 7.65 (dd, *J* = 2.0, 8.4 Hz, 1H), 7.51 (d, *J* = 1.6 Hz, 1H), 6.24 (s, 1H), 6.01 (s, 1H), 3.96 (s, 3H).

#### Methyl 4-bromo-2-(1r,2r)-1-cyano-2-fluorocyclopropyl)benzoate and methyl 4-bromo-2-(1r,2s)-1-cyano-2-fluorocyclopropyl)benzoate

To a solution of methyl 4-bromo-2-(1-cyanovinyl)benzoate (800 mg, 3.01 mmol) in THF (30 mL) at 0 °C were added (fluoromethyl)(phenyl)(2,3,4,5-tetramethylphenyl)sulfonium tetrafluoroborate (2.3 g, 6.0 mmol, **Intermediate 1**) and NaH (1.2 g, 30.1 mmol, 60% purity in mineral oil). The reaction mixture was stirred at 20 °C for 1 h. The reaction mixture was poured into water (50 mL) and extracted with EtOAc (3 × 30 mL). The combined organic layers were washed with brine (10 mL), dried over anhydrous Na_2_SO_4_, filtered, and concentrated under reduced pressure. The residue was purified by silica gel column chromatography to provide the title products as a 3:2 mixture. LCMS: *m/z* = 298.0, 300.0 [M+H]^+^.

#### (2’r,4s)-6-bromo-2’-fluorospiro[2,3-dihydroisoquinoline-4,1’-cyclopropane]-1-one and (2’r,4r)-6-bromo-2’-fluorospiro[2,3-dihydroisoquinoline-4,1’-cyclopropane]-1-one

To a solution of methyl 4-bromo-2-(1r,2r)-1-cyano-2-fluorocyclopropyl)benzoate and methyl 4-bromo-2-(1r,2s)-1-cyano-2-fluorocyclopropyl)benzoate (0.92 g, 3.09 mmol, 3:2 mixture) and CoCl_2_ (401 mg, 3.09 mmol) in MeOH (10 mL) at 0 °C was added NaBH_4_ (350 mg, 9.26 mmol). The reaction mixture was stirred at 20 °C for 2 h. The reaction mixture was poured into sat. aq. NH_4_Cl (20 mL) and extracted with EtOAc (3 × 20 mL). The combined organic layers were washed with brine (2 × 10 mL), dried over anhydrous Na_2_SO_4_, filtered, and concentrated under reduced pressure. The residue was purified by silica gel column chromatography to provide:

(2’r,4s)-6-bromo-2’-fluorospiro[2,3-dihydroisoquinoline-4,1’-cyclopropane]-1-one: ^1^H NMR (400 MHz, CDCl_3_): δ 8.02 (d, *J* = 8.4 Hz, 1H), 7.52 (dd, *J* = 2.0, 8.4 Hz, 1H), 6.85 (d, *J* = 2.0 Hz, 1H), 6.12 (br s, 1H), 4.70-4.50 (m, 1H), 3.87 (d, *J* = 12.4 Hz, 1H), 3.51 (dd, *J* = 4.4, 12.8 Hz, 1H), 1.62-1.54 (m, 1H), 1.42-1.24 (m, 1H).

(2’r,4r)-6-bromo-2’-fluorospiro[2,3-dihydroisoquinoline-4,1’-cyclopropane]-1-one: ^1^H NMR (400 MHz, CDCl_3_): δ 8.02 (d, *J* = 8.4 Hz, 1H), 7.55 (dd, *J* = 2.0, 8.4 Hz, 1H), 7.28 (d, *J* = 2.0 Hz, 1H), 6.56 (br s, 1H), 4.81-4.59 (m, 1H), 3.78 (dd, *J* = 7.6, 12.4 Hz, 1H), 2.74 (dd, *J* = 4.4, 12.8 Hz, 1H), 1.83-1.73 (m, 1H), 1.21-1.15 (m, 1H).

#### Methyl 2-[(2’r,4s)-6-bromo-2’-fluoro-1-oxospiro[3H-isoquinoline-4,1’-cyclopropane]-2-yl]acetate

To a solution of (2’r,4s)-6-bromo-2’-fluorospiro[2,3-dihydroisoquinoline-4,1’-cyclopropane]-1-one (200 mg, 0.74 mmol) in DMF (2.0 mL) at 0 °C was added NaH (45 mg, 1.11 mmol, 60% purity in mineral oil). The reaction mixture was stirred at 20 °C for 0.5 h followed by the addition of methyl 2-bromoacetate (227 mg, 1.48 mmol). The reaction mixture was stirred at 20 °C for 3 h. The reaction mixture was poured into sat. aq. NH_4_Cl and ice water mixture (10 mL) and extracted with EtOAc (3 × 5mL). The combined organic layers were washed with brine (5 mL), dried over anhydrous Na_2_SO_4_, filtered, and concentrated under reduced pressure. The residue was purified by silica gel column chromatography. The residue was further purified by chiral SFC: (Column: Daicel Chiralpak AD (250mm x 30mm, 10 µm particle size); Mobile Phase: A: CO_2_, B: 0.1% NH_4_OH in *i*-PrOH; Gradient: 38%B isocratic; Flow Rate: 64 g/min; Detection Wavelength: 220 nm; Column Temperature: 40 °C; System Back Pressure: 100 bar) to provide:

methyl 2-[(2’s,4r)-6-bromo-2’-fluoro-1-oxospiro[3H-isoquinoline-4,1’-cyclopropane]-2-yl]acetate (first eluting isomer): LCMS: *m/z* = 342.0, 344.0 [M+H]^+^. ^1^H NMR (400 MHz, CDCl_3_): δ 8.02 (d, *J* = 8.0 Hz, 1H), 7.51 (dd, *J* = 1.6, 8.4 Hz, 1H), 6.83 (d, *J* = 1.6 Hz, 1H), 4.73-4.54 (m, 2H), 4.10 (dd, *J* = 2.0, 12.4 Hz, 1H), 4.00 (d, *J* = 17.2 Hz, 1H), 3.77 (s, 3H), 3.44 (d, *J* = 12.4 Hz, 1H), 1.60-1.57 (m, 1H), 1.44-1.33 (m, 1H).

methyl 2-[(2’s,4r)-6-bromo-2’-fluoro-1-oxospiro[3H-isoquinoline-4,1’-cyclopropane]-2-yl]acetate (second eluting isomer): LCMS: *m/z* = 342.0, 344.0 [M+H]^+^. ^1^H NMR (400 MHz, CDCl_3_): δ 8.02 (d, *J* = 8.0 Hz, 1H), 7.51 (dd, *J* = 1.6, 8.4 Hz, 1H), 6.83 (d, *J* = 1.6 Hz, 1H), 4.73-4.54 (m, 2H), 4.10 (dd, *J* = 2.0, 12.4 Hz, 1H), 4.00 (d, *J* = 17.2 Hz, 1H), 3.77 (s, 3H), 3.44 (d, *J* = 12.4 Hz, 1H), 1.60-1.57 (m, 1H), 1.44-1.33 (m, 1H).

#### Methyl 2-(2’r,4s)-2-fluoro-6’-iodo-1’-oxo-1’*H*-spiro[cyclopropane-1,4’-isoquinolin]-2’(3’*H*)-yl)acetate

To a solution of methyl 2-[(2’r,4s)-6-bromo-2’-fluoro-1-oxospiro[3H-isoquinoline-4,1’-cyclopropane]-2-yl]acetate (6.0 g, 11.5 mmol, second eluting isomer) in toluene (180 mL) were added NaI (10.5 g, 70.1 mmol), *N*,*N*’-dimethyl-1,2-diaminocyclohexane (998 mg, 7.01 mmol) and CuI (668 mg, 3.51 mmol). The mixture was stirred at 130 °C for 72 h. The mixture was poured into a mixture of H_2_O (200 mL) and EtOAc (80 mL). The mixture was filtered, and the filtrate was extracted with EtOAc (2 × 80 mL). The combined organic layers were washed with brine (80 mL), dried over anhydrous Na_2_SO_4_, filtered and concentrated under reduced pressure. The residue was purified by silica gel column chromatography. LCMS: *m/z*: 389.9 [M+H]^+^. ^1^H NMR (400 MHz, CDCl_3_): δ 7.86 (d, *J* = 8.4 Hz, 1H), 7.73 (dd, *J* = 1.6, 8.4 Hz, 1H), 7.04 (d, *J* = 1.6 Hz, 1H), 4.76-4.47 (m, 1H), 4.09 (dd, *J* = 2.0, 12.4 Hz, 1H), 3.99 (d, *J* = 17.2 Hz, 1H), 3.77 (s, 3H), 3.43 (d, *J* = 12.8 Hz, 1H), 1.62-1.59 (m, 1H), 1.58-1.54 (m, 1H), 1.43-1.32 (m, 1H).

#### 2-(2’r,4s)-2-fluoro-6’-iodo-1’-oxo-1’*H*-spiro[cyclopropane-1,4’-isoquinolin]-2’(3’*H*)-yl)acetic acid

To a solution of methyl 2-(2’r,4s)-2-fluoro-6’-iodo-1’-oxo-1’*H*-spiro[cyclopropane-1,4’-isoquinolin]-2’(3’*H*)-yl)acetate (6.0 g, 15.4 mmol) in THF (120 mL) and H_2_O (60 mL) was added LiOH·H_2_O (2.59 g, 61.67 mmol). The mixture was stirred for 12 h. The mixture was diluted with H_2_O (200 mL) and extracted with MTBE (150 mL). The aqueous phase was separated, and the pH was adjusted to pH = 3∼4 by addition of 3 N HCl. The mixture was extracted with EtOAc (3 × 150 mL). The combined organic layers were washed with brine (150 mL), dried over anhydrous Na_2_SO_4_, filtered and concentrated under reduced pressure. The crude residue was used directly. LCMS: *m/z*: 375.8 [M+H]^+^. ^1^H NMR (400 MHz, DMSO-d_6_): δ 12.78 (s, 1H), 7.77 (d, *J* = 8.0 Hz, 1H), 7.65 (d, *J* = 8.0 Hz, 1H), 7.36 (s, 1H), 5.22-4.86 (m, 1H), 4.35 (d, *J* = 17.2 Hz, 1H), 4.04 (d, *J* = 17.2 Hz, 1H), 3.87 (d, *J* = 12.8 Hz, 1H), 3.51 (d, *J* = 13.2 Hz, 1H), 1.83-1.62 (m, 1H), 1.53-1.38 (m, 1H).

#### 2-[(2’r,4s)-2’-fluoro-6-iodo-1-oxospiro[3H-isoquinoline-4,1’-cyclopropane]-2-yl]-N-(3-hydroxy-3-methylcyclobutyl)acetamide (TN-101)

To a mixture of 2-(2’r,4s)-2-fluoro-6’-iodo-1’-oxo-1’*H*-spiro[cyclopropane-1,4’-isoquinolin]-2’(3’*H*)-yl)acetic acid (4.1 g, 10.9 mmol) and (1*s*,3*s*)-3-amino-1-methylcyclobutanol hydrochloride (1.80 g, 13.1 mmol) in DCM (40 mL) was added DIEA (5.65 g, 43.7 mmol), EDCI (2.51 g, 13.1 mmol) and HOBt (1.77 g, 13.12 mmol). The mixture was stirred at 30 °C for 12 h. The mixture was poured into H_2_O (150 mL) and EtOAc (150 mL). The mixture was filtered, and the filtrate was separated. The organic layer was washed with aq. sat. NaHCO_3_, dried over anhydrous Na_2_SO_4_, filtered and concentrated under reduced pressure. The above procedure was repeated 5 times to acquire 18.6 g of crude residue. The combined residue was triturated with MeCN (150 mL) and dried under reduced pressure. The residue was then recrystallized: the residue was suspended in EtOAc (400 mL) and heated to 100 °C with stirring. EtOAc (16 × 100 mL) was added to the solution at 100 °C in portions until the solid was mostly dissolved. The mixture was filtered, and the filtrate continued to be stirred at 100 °C. EtOAc (2.5 × 100 mL) was added to the solution at 100 °C in portions until the solid was dissolved. Then n-heptane (12 × 100 mL) was added to the solution at 100 °C in portions, until the solution was cloudy. The mixture was cooled to 80 °C for 30 mins and stirred for another 10 mins. The mixture was cooled to 50 °C for 30 mins and stirred for another 10 mins. The mixture was cooled to 40 °C for 30 mins and stirred for another 10 mins. The mixture was cooled to 20 °C and stirred for 12 h. The mixture was filtered, and the filter cake was washed with heptane. The filter cake was dried under reduced pressure.

**LCMS**: *m/z* = 459.1 [M+H]^+^. (99.4 % purity)

**Chiral SFC**: 99.9 % e.e.

**^1^H NMR**: (400MHz, DMSO-d_6_): δ 8.16 (d, *J* = 7.2 Hz, 1H), 7.76 (dd, *J* = 1.2, 8.4 Hz, 1H), 7.64 (d, *J* = 8.4 Hz, 1H), 7.34 (s, 1H), 5.15-4.86 (m, 2H), 4.29 (d, *J* = 16.4 Hz, 1H), 3.93-3.80 (m, 2H), 3.80-3.69 (m, 1H), 3.44 (d, *J* = 13.2 Hz, 1H), 2.21 (dt, *J* = 4.0, 7.2 Hz, 2H), 1.93 (t, *J* = 10.0 Hz, 2H), 1.70 (td, *J* = 7.2, 12.0 Hz, 1H), 1.54-1.38 (m, 1H), 1.21 (s, 3H).

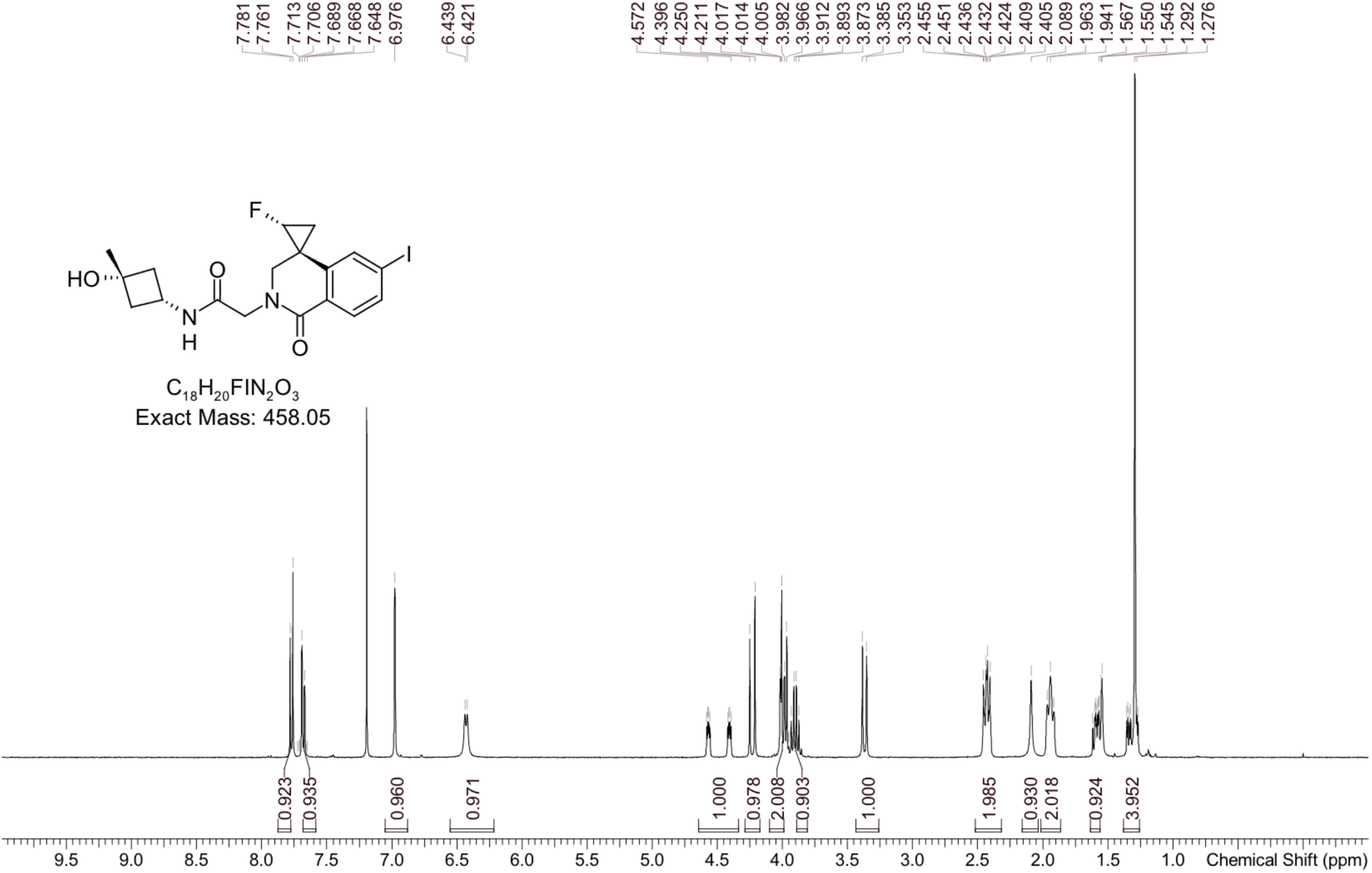

#### Polarizing light microscope report

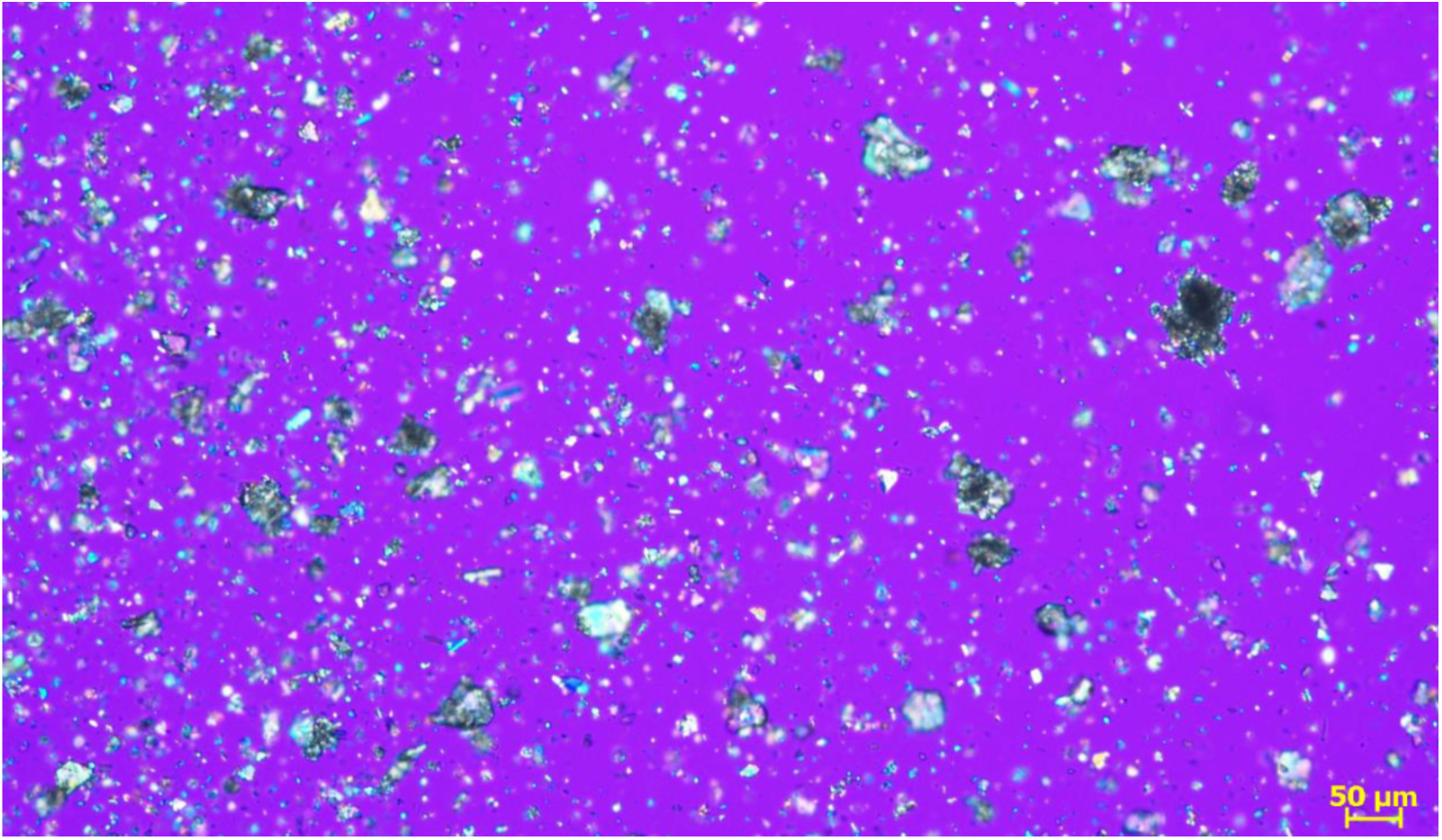

#### XRPD

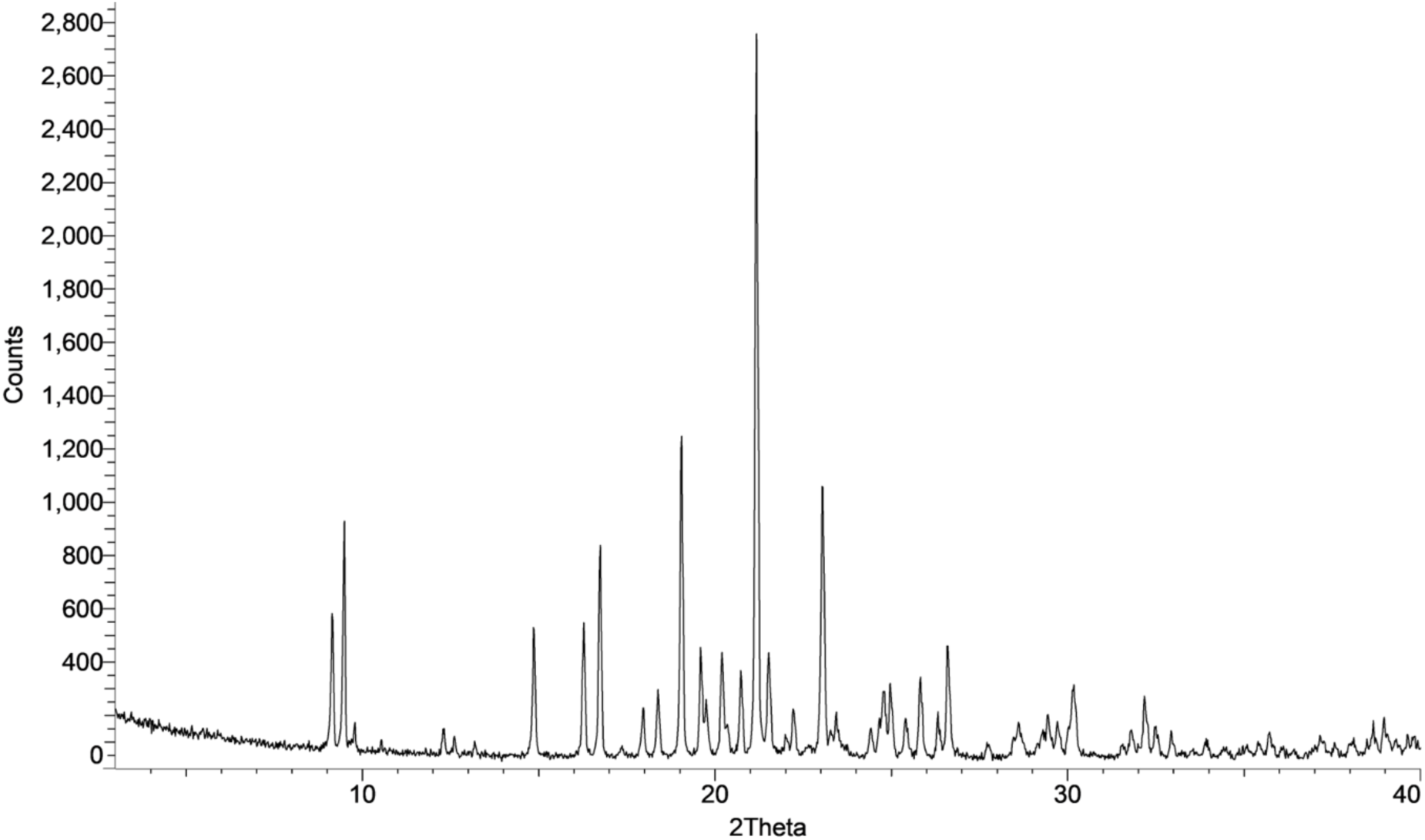

#### DSC

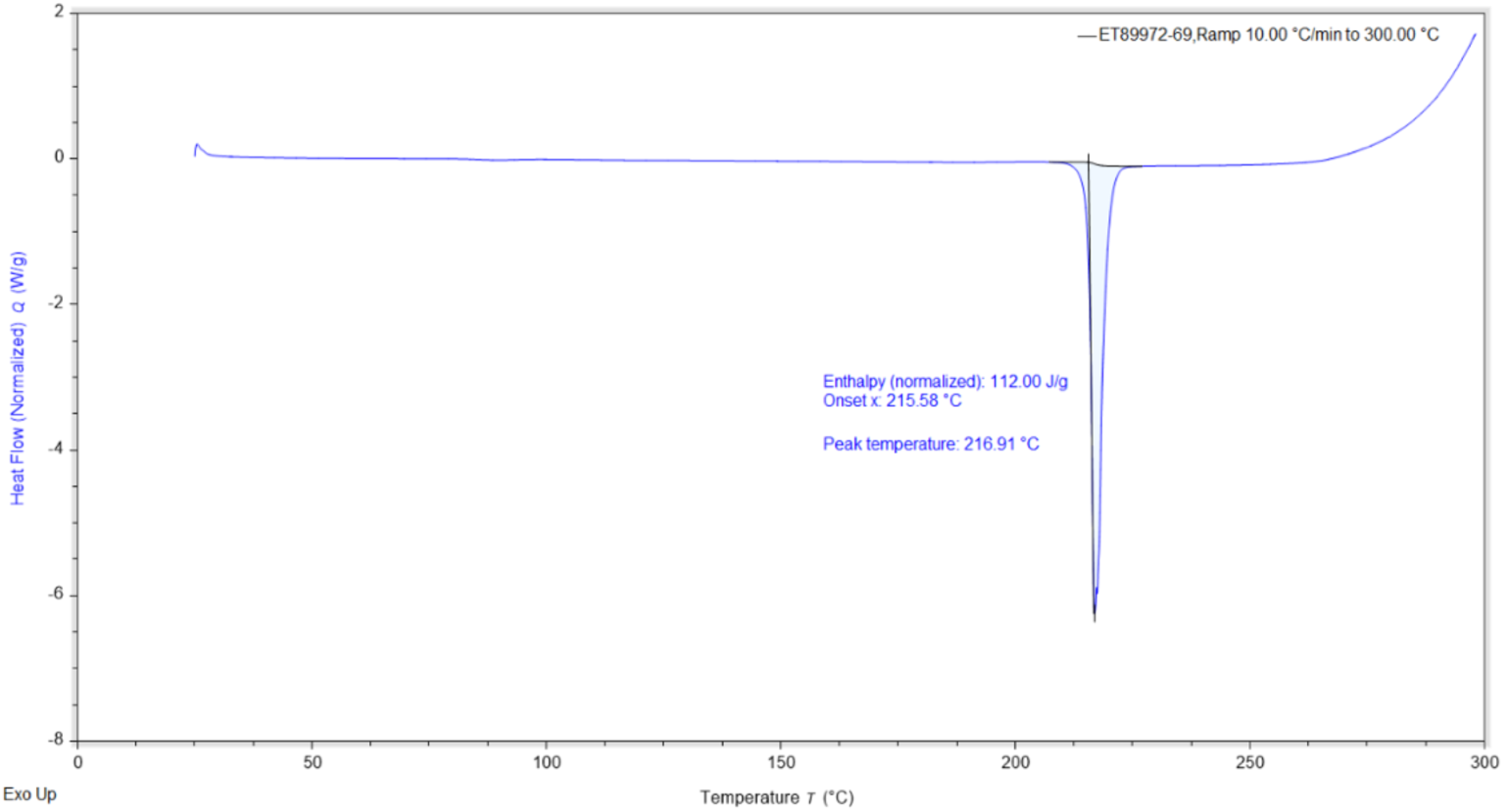

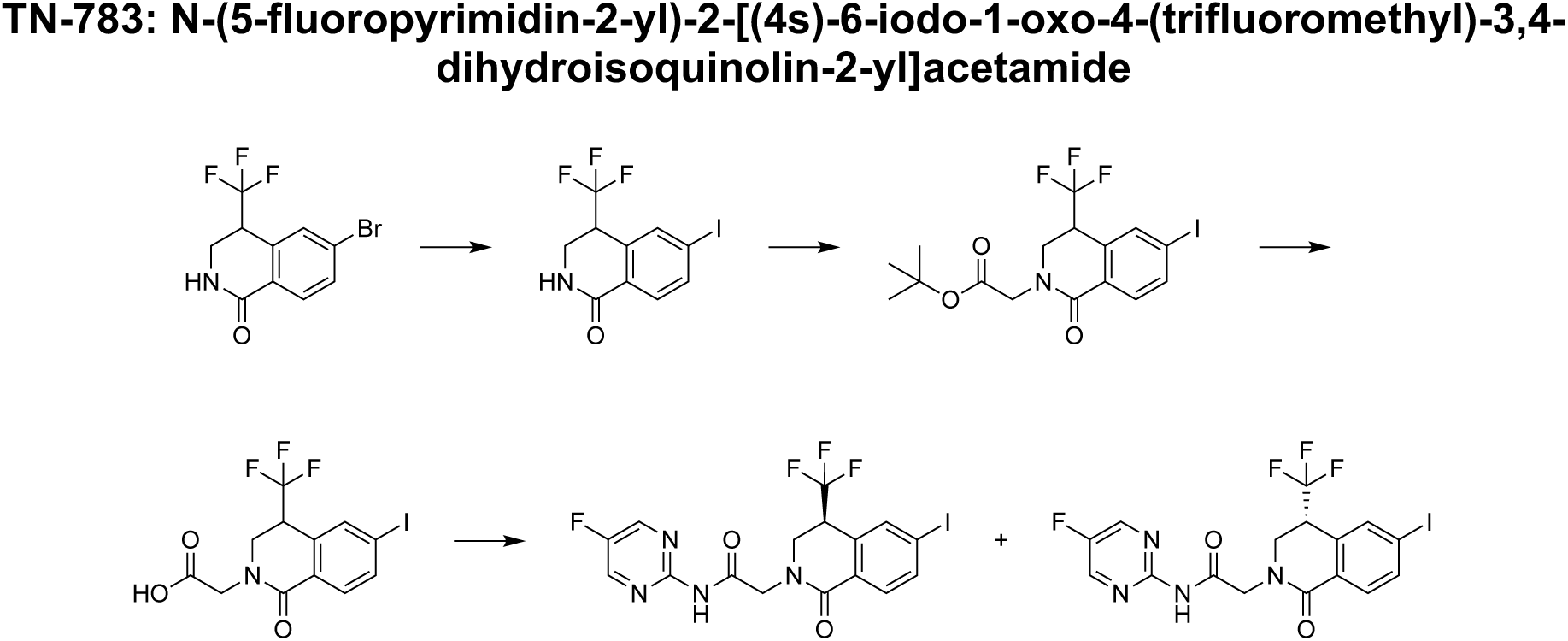

#### 6-bromo-4-(trifluoromethyl)-3,4-dihydro-2H-isoquinolin-1-one

Can be prepared according to the procedures outlined in WO2022/232632.

#### 6-iodo-4-(trifluoromethyl)-3,4-dihydro-2H-isoquinolin-1-one

To a solution of 6-bromo-4-(trifluoromethyl)-3,4-dihydro-2H-isoquinolin-1-one (100 g, 340 mmol) and N,N’-dimethylethane-1,2-diamine (15.0 g, 170 mmol) in 1,4-dioxane (1.0 L) were added NaI (152.0 g, 1.02 mol) and CuI (64.7 g, 340 mmol). The reaction mixture was stirred at 125°C for 10 h. The suspension was filtered through a pad of Celite and the filter cake was washed with EtOAc (2 x 300 mL). The combined filtrates were diluted with H_2_O (2.0 L) and extracted with EtOAc (2 x 300 mL). The combined organic layers were washed with brine, dried over anhydrous Na_2_SO_4_, filtered and concentrated under reduced pressure. The residue was triturated with petroleum ether/EtOAc (5:1, 2.0 L) for 1 h. The mixture was filtered, and the filter cake was concentrated under reduced pressure. ^1^H NMR (400 MHz, CDCl_3_): δ 7.95-7.84 (m, 2H), 7.77 (s, 1H), 6.25 (br s, 1H), 3.95-3.79 (m, 2H), 3.61-3.44 (m, 1H).

#### tert-butyl 2-[6-iodo-1-oxo-4-(trifluoromethyl)-3,4-dihydroisoquinolin-2-yl]acetate

To a solution of 6-iodo-4-(trifluoromethyl)-3,4-dihydro-2H-isoquinolin-1-one (200 g, 586 mmol) in THF (1.6 L) at -20 °C was added NaH (60% purity, 28.1 g, 703 mmol). The mixture was stirred at -20 °C for 5 min. A solution of tert-butyl 2-bromoacetate (137 g, 703 mmol) in THF (400 mL) was added dropwise at -20 °C. The reaction was stirred at 0 °C for 4 h. The reaction mixture was poured in aq. sat. NH_4_Cl (3.0 L) and extracted with MTBE (3 x 2.0 L). The combined organic layers were washed with brine, dried over anhydrous Na_2_SO_4_, filtered and concentrated under reduced pressure. The residue was filtered through a silica gel pad using EtOAc as the eluent and concentrated under reduced pressure. The residue was triturated with petroleum ether/MTBE (4:1, 2.0 L) for 30 min. The mixture was filtered and the filter cake was concentrated under reduced pressure. ^1^H NMR (400 MHz, CDCl_3_): δ 7.93-7.84 (m, 2H), 7.73 (s, 1H), 4.73 (d, J = 17.2 Hz, 1H), 4.15 (ddd, J = 0.8, 4.8, 13.2 Hz, 1H), 3.80-3.67 (m, 2H), 3.58-3.51 (m, 1H), 1.48 (s, 9H).

#### 2-[6-iodo-1-oxo-4-(trifluoromethyl)-3,4-dihydroisoquinolin-2-yl]acetic acid

To a solution of tert-butyl 2-[6-iodo-1-oxo-4-(trifluoromethyl)-3,4-dihydroisoquinolin-2-yl]acetate (250 g, 550 mmol) in 1,4-dioxane (250 mL) at 0 °C was added HCl/dioxane (2.25 L, 4M). The reaction mixture was stirred at 45 °C for 5 h. The reaction was concentrated under reduced pressure. The residue was triturated with petroleum ether/MTBE (5:1, 250 mL) for 30 min. The mixture was filtered and the filter cake was concentrated under reduced pressure. ^1^H NMR (400 MHz, CDCl_3_): δ 7.95 (dd, J = 1.6, 8.0 Hz, 1H), 7.90 (s, 1H), 7.71 (d, J = 8.0 Hz, 1H), 4.44 (d, J = 17.6 Hz, 1H), 4.26-4.10 (m, 1H), 4.07-3.96 (m, 1H), 3.86 (d, J = 13.6 Hz, 1H), 3.56 (s, 2H).

#### N-(5-fluoropyrimidin-2-yl)-2-[(4s)-6-iodo-1-oxo-4-(trifluoromethyl)-3,4-dihydroisoquinolin-2-yl]acetamide (TN-783)

To a solution of 2-[6-iodo-1-oxo-4-(trifluoromethyl)-3,4-dihydroisoquinolin-2-yl]acetic acid (120 g, 300 mmol) and 5-fluoropyrimidin-2-amine (68 g, 601 mmol) in MeCN (1.8 L) were added 2-chloro-1-methylpyridinium iodide (92 g, 360 mmol) and pyridine (71.3 g, 902 mmol). The reaction mixture was stirred at 60 °C for 12 h. The mixture was concentrated under reduced pressure. The residue was dissolved in DCM and washed with aq. HCl (2 x 1.5 L, 1M). The combined organic layers were washed with aq. sat. NaHCO_3_ (2.0 L) and brine, dried over anhydrous Na_2_SO_4_, filtered and concentrated under reduced pressure. The residue was triturated with MeCN (4.0 L) for 30 min. The residue was further purified by chiral SFC column: DAICEL CHIRALCEL OZ 250 × 50 mm I.D. 10μm particle size; mobile phase: A: CO2, B: EtOH; Gradient: 50% B isocratic elution mode, detection wavelength: 220 nm; column temperature: 40 °C; system back pressure: 100 bar) to provide:

#### N-(5-fluoropyrimidin-2-yl)-2-[(4s)-6-iodo-1-oxo-4-(trifluoromethyl)-3,4-dihydroisoquinolin-2-yl]acetamide (TN-783, first eluting isomer)

LCMS: *m/z* **=** 495.0 [M+H]^+^. ^1^H NMR (400 MHz, CDCl_3_): δ 10.98 (s, 1H), 8.77 (s, 2H), 7.96 (d, J = 8.4 Hz, 1H), 7.91 (s, 1H), 7.71 (d, J = 8.0 Hz, 1H), 4.89 (br d, J = 16.8 Hz, 1H), 4.30-4.14 (m, 2H), 4.13-4.03 (m, 1H), 3.88 (br d, J = 14.0 Hz, 1H). (99.5 % purity, 99.48% e.e.)&

#### N-(5-fluoropyrimidin-2-yl)-2-[(4r)-6-iodo-1-oxo-4-(trifluoromethyl)-3,4-dihydroisoquinolin-2-yl]acetamide (Second eluting isomer)

LCMS: *m/z* = 495.0 [M+H]^+^. ^1^H NMR (400 MHz, CDCl_3_): δ 10.98 (s, 1H), 8.77 (s, 2H), 7.96 (d, J = 8.4 Hz, 1H), 7.91 (s, 1H), 7.71 (d, J = 8.0 Hz, 1H), 4.89 (br d, J = 16.8 Hz, 1H), 4.30-4.14 (m, 2H), 4.13-4.03 (m, 1H), 3.88 (br d, J = 14.0 Hz, 1H).

N-(5-fluoropyrimidin-2-yl)-2-[(4s)-6-iodo-1-oxo-4-(trifluoromethyl)-3,4-dihydroisoquinolin-2-yl]acetamide (TN-783, first eluting isomer, 150 g) was suspended in EtOH (450 mL) and stirred for 30 min. H_2_O (3 x 500 mL) was added to the solution in portions for 30 min and the mixture was stirred for an additional 30 min. The mixture was filtered and the filter cake was washed with H_2_O (3 x 500 mL) and concentrated under reduced pressure. The resulting solid was dried in a vacuum drying oven at 50 °C for 12 h to afford crystalline N-(5-fluoropyrimidin-2-yl)-2-[(4s)-6-iodo-1-oxo-4-(trifluoromethyl)-3,4-dihydroisoquinolin-2-yl]acetamide (TN-783).

**LCMS**: *m/z* = 495.0 [M+H]^+^. (99.5 % purity)

**Chiral SFC**: 99.48% e.e.

**^1^H NMR**: (400 MHz, CDCl_3_): δ 10.98 (s, 1H), 8.77 (s, 2H), 7.96 (d, J = 8.4 Hz, 1H), 7.91 (s, 1H), 7.71 (d, J = 8.0 Hz, 1H), 4.89 (br d, J = 16.8 Hz, 1H), 4.30-4.14 (m, 2H), 4.13-4.03 (m, 1H), 3.88 (br d, J = 14.0 Hz, 1H).

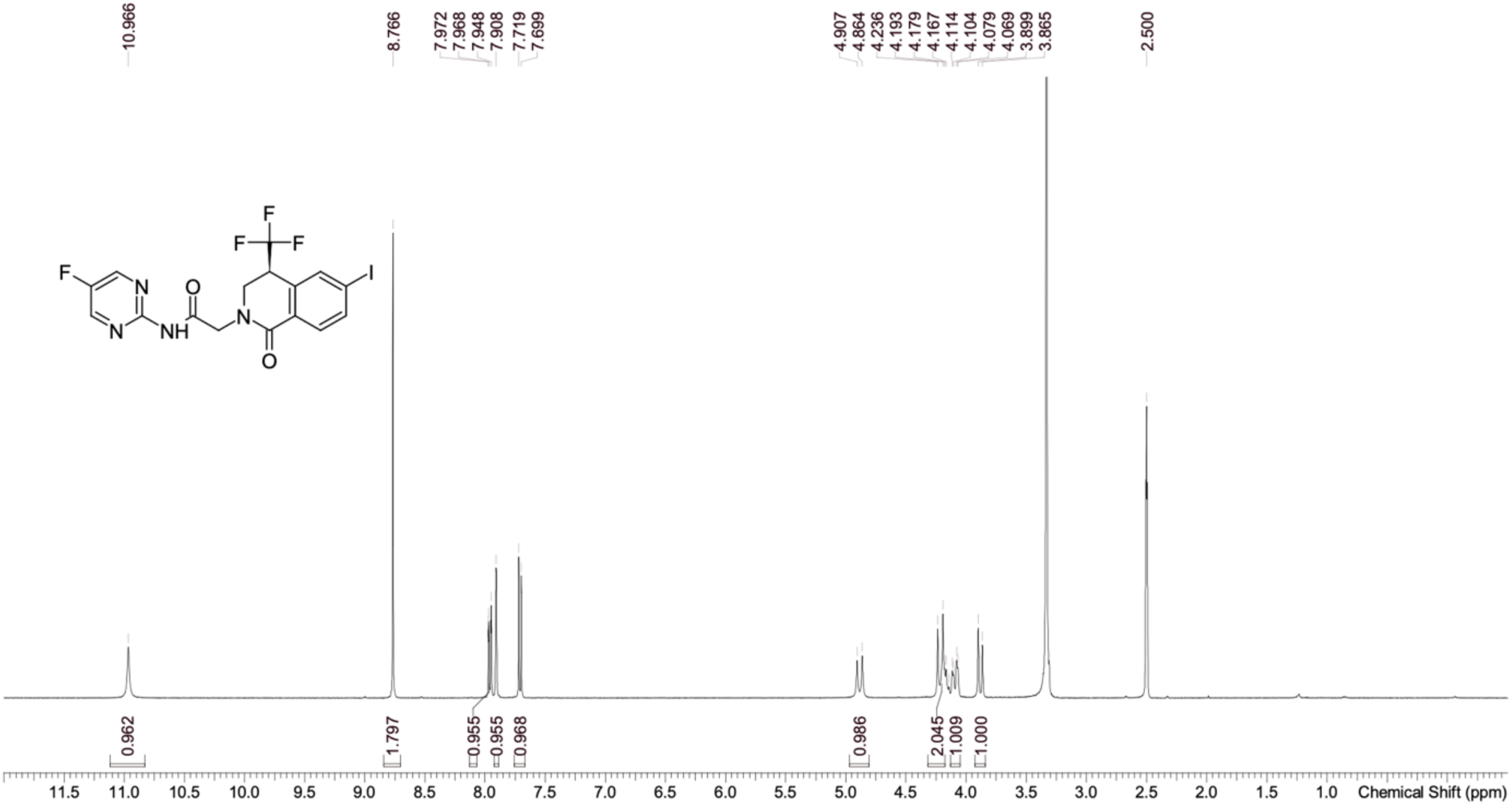

#### Polarizing Light Microscope Report

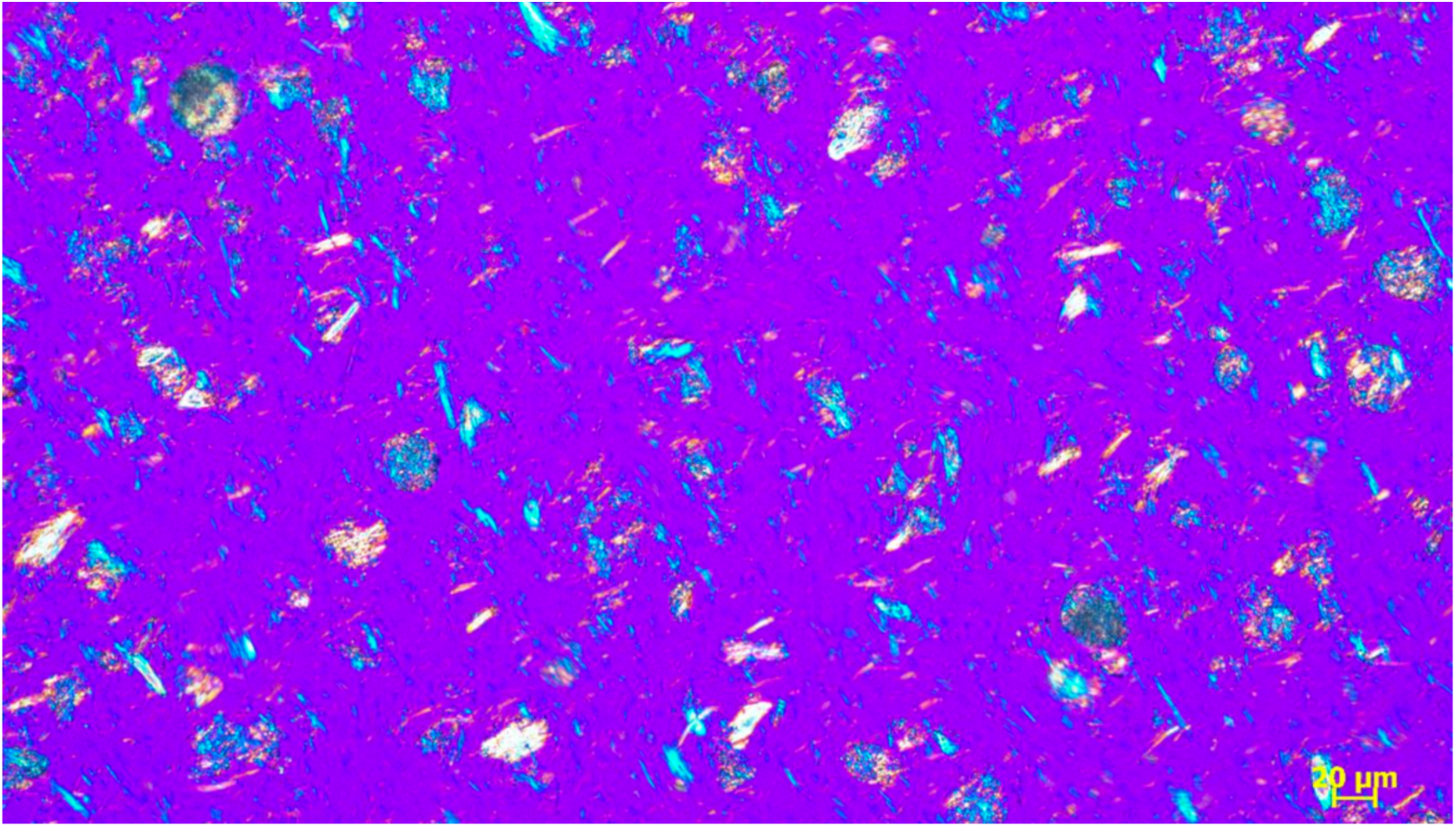

#### XRPD

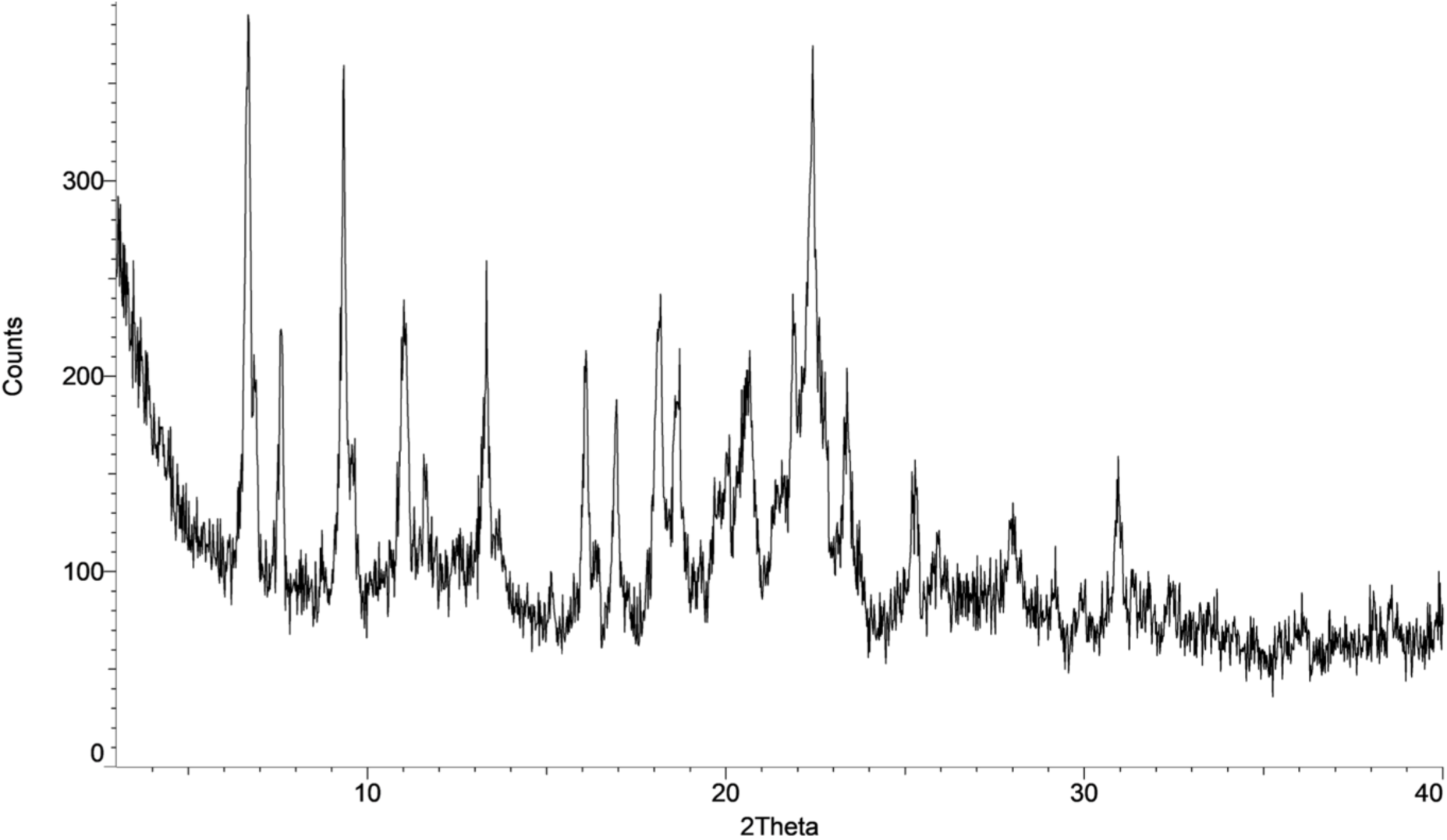

#### DSC

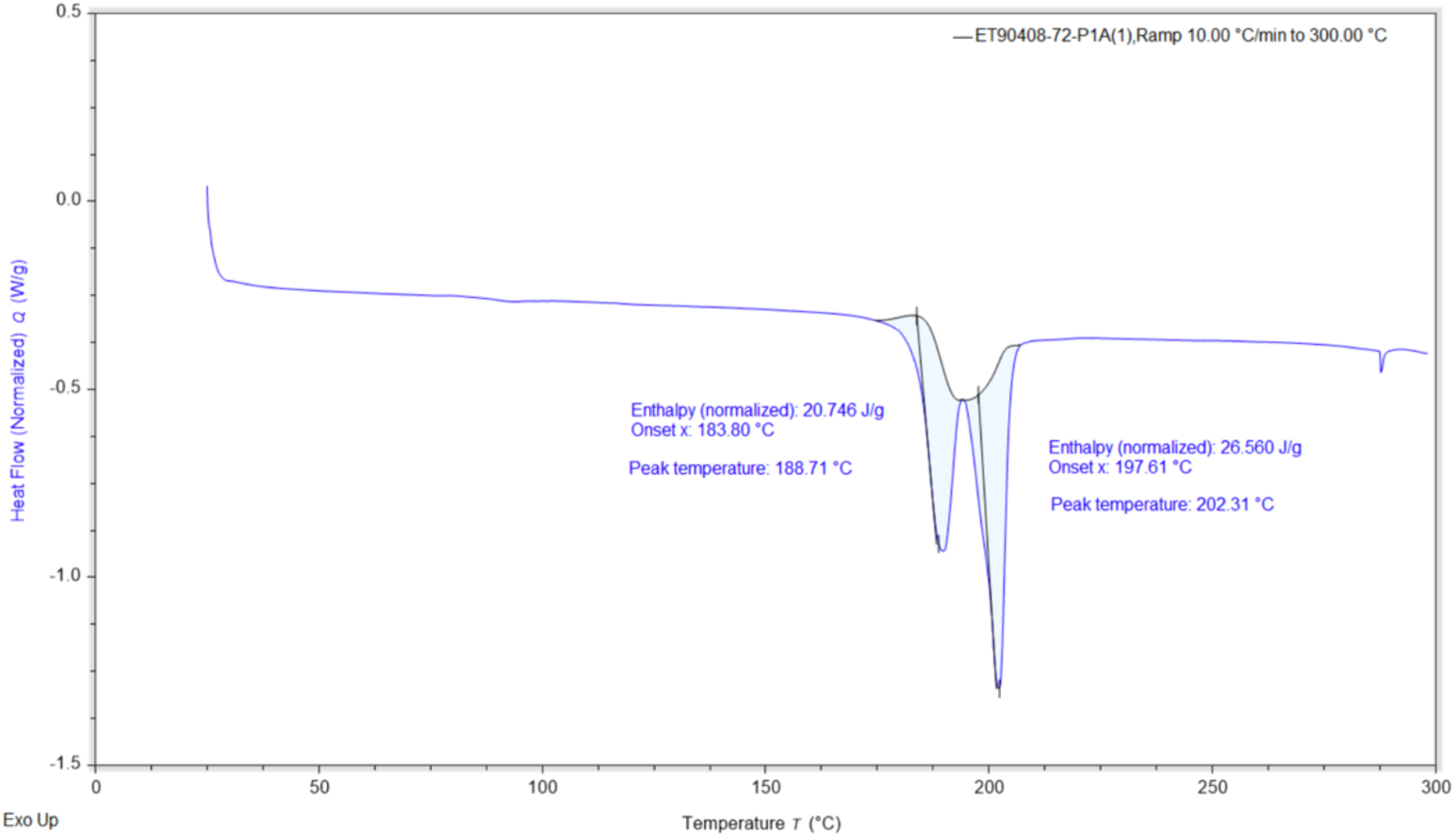

### Protein expression and purification

Full-length human NEK7 (residues 1–302) was cloned into a pET28a vector with an N-terminal 6×His–SUMO tag and expressed in E. coli BL21(DE3) cells. Cultures were grown in LB medium containing 50 µg/mL kanamycin and protein expression induced overnight at 15 °C with 0.5 mM IPTG. Cells were harvested and lysed in buffer (50 mM Tris-HCl pH 7.5, 500 mM NaCl, 5 mM MgCl₂, 10 mM imidazole, 10% glycerol, 2 mM β-mercaptoethanol). NEK7 was purified by Ni-affinity chromatography, followed by Ulp1 protease cleavage of the SUMO tag and a second Ni-affinity step. Final purification was performed by size-exclusion chromatography in 20 mM Tris-HCl pH 7.5, 150 mM NaCl, and 0.5 mM TCEP.

Human NLRP3 (residues 136–1034; R137C, K138A, K142A) was cloned into a pFastBac1 vector with an N-terminal MBP tag and expressed in Hi-5 insect cells for 48 h post-infection at 27 °C. Cells were lysed in buffer (50 mM Tris-HCl pH 7.5, 500 mM NaCl, 0.5 mM TCEP, 10% glycerol, protease inhibitor cocktail) and clarified lysate applied to a Dextrin Sepharose HP column. MBP-tagged protein was eluted with 25 mM maltose and further purified by size-exclusion chromatography in 20 mM Tris-HCl pH 7.5, 150 mM NaCl, 0.5 mM TCEP, and 10% glycerol.

### NLRP3/NEK7 cryo-EM structure determination

The NLRP3–NEK7 complex was assembled by overnight incubation at 4 °C, further purified by size-exclusion chromatography, and incubated with 0.3 mM ADP, 5 mM MgCl₂, and 50 μM TN-551 prior to vitrification (76). Purified complex (0.45 mg/mL) was applied to glow-discharged Quantifoil R1.2/1.3 300 mesh copper grids, blotted for 5 s at 4 °C and 100% humidity, and plunge-frozen in liquid ethane using a Vitrobot Mark IV (76).

High-resolution cryo-EM data were collected on a 300 kV Titan Krios equipped with a Gatan K3 direct electron detector in super-resolution counting mode. A total of 5562 movies were recorded with SerialEM at a physical pixel size of 0.83 Å, a defocus range of −1.0 to −2.5 µm, and a total electron dose of ∼81 e⁻/Å² fractionated over 50 frames, using collection strategies similar to those previously reported for NLRP3–NEK7 complexes (76).

Movies were motion-corrected and CTF parameters estimated using RELION 3.0 and cryoSPARC v3.0.1 (76). Automated particle picking followed by iterative 2D and 3D classification yielded 340,724 monomer particles and 194,515 dimer particles, which were refined to maps at 2.61 Å and 3.02 Å resolution, respectively (FSC 0.143). Local refinements improved density quality, enabling the modeling of additional NEK7 residues (aa 28–112) not resolved in earlier reconstructions. Atomic models were built in Chimera and Coot and refined using CCPEM, consistent with established workflows (76).

### NLRP3 fluorescence polarization displacement assay

An assay similar to that developed by Dekker et al. was used to measure compound binding to purified NLRP3 via fluorescent tracer displacement (25). Test compounds were spotted on a black 384 well plate (Thermo Fisher, 262260) by an Echo acoustic dispenser. Purified NLRP3 protein was diluted to 0.0625 µM in fluorescence polarization buffer (20 mM HEPES, 130 mM sodium gluconate, 5 mM NaHCO3, 20 mM magnesium gluconate, 3 mM potassium gluconate, 100 mM DL-serine (Sigma, AlS4375), 0.005% (wt/vol) Tween 20 (Sigma, PI379), 0.05% (wt/vol) PEI (Sigma, 764604) with 100 uM ADP (Sigma, A2754). 20 uL of the NLRP3/ADP mix was added to the plate using a Multidrop liquid dispenser. A fluorescently labeled MCC950 analogue was dispensed on the plate at 0.0625 µM final concentration using TECAN D300e. The plate was incubated at room temperature shaking at 500 rpm for 60 minutes. The plate was read by Envision plate reader using FITC FP filters. Data was normalized to in-plate controls: no-treatment (DMSO) and positive control compound.

### THP-1 LPS/nigericin stimulation assay

THP-1 (ATCC, TIB-202) cells were cultured in RPMI 1640 medium (Gibco, 61870036) supplemented with 10% Hyclone FBS (Cytiva, SH3008003) and 1x Pen Strep (Gibco,15140122). One day before treatment, cells were plated and differentiated by adding 25 ng/mL IFN-γ (PeproTech, 300-02-100UG) to 100,000 cells in 100 uL culture medium (RPMI 1640, 10% FBS, 1% pen/strep, 0.05 mM 2-mercaptoethanol) and dispensing into a 96 well tissue-culture treated plate. After 24 hours the IFN-γ containing media was removed and replaced with 100 uL culture media containing 50 ng/mL LPS (Invivogen, thrl-3pelps) and incubated for 4 hours. Compounds dissolved in DMSO were then spotted using Tecan D300e and left to incubate for 1 hour. 20 uL ATP was added to a final concentration of 5 mM and incubated for 1 hour. The supernatant was collected and stored at -20 °C until tested.

ELISA plates (Sigma, p6366) were coated with capture antibody (MT175) in PBS at 2 ug/mL overnight at 4 C. Capture antibody was removed and the plate was washed 4 times with PBST. The plate was blocked with 25 uL/well blocking buffer (LiCor, 927-70001) containing 0.1% Tween 20 for 1 hour at room temperature. Following blocking the plate was washed 4 times with PBST. 25 uL/well of test sample was added and incubated for 2 hours at room temperature. The plate was washed 4 times with PBST and then incubated with 15 uL/well mAb7P10-biotin at 0.5 ug/mL in blocking buffer for 1 hour. The antibody was then removed, and the plate washed 4 times with PBST. 15 uL/well streptavidin-HRP (1:1000) in blocking buffer was added and incubated for 1 hour followed by washing the plate 4 times in PBST. HRP substrate was added (25 uL/well) and incubated 1 minute at room temperature, followed by the addition of 25 uL/well stop solution. The plate absorbance was read at 450 nm. Data was normalized by in-plate untreated (DMSO) and positive control compound treated conditions.

### THP-1 palmitate challenge assay

THP-1 cells were seeded at a density of 7.5 × 10^4^ cells per well onto a poly-D-lysine (PDL)-coated 96-well plate (Corning, 354640) and differentiated into macrophage-like cells using 100 nM phorbol 12-myristate 13-acetate (PMA; Sigma-Aldrich, P1585) for 24 hours prior to treatment. After differentiation, the culture medium was replaced with serum-free RPMI 1640 medium. Cells were pre-incubated with a 10-point titration of NLRP3 inhibitors (0.3 nM–6 µM) for 1 h at 37 °C, with each concentration tested in technical duplicates. Following pre-incubation, cells were stimulated overnight at 37 °C with palmitate-BSA complex (Cayman Chemical, 29558) at a final concentration of 400 µM palmitate and 64 µM BSA. Conditioned media were collected and transferred to a 96-well V-bottom plate (Greiner, 651201), then centrifuged at 300 × g for 5 min. Supernatants were transferred to PCR tubes (Thermo Fisher, AB2005), snap-frozen on dry ice, and stored at -80 °C until analysis. IL-1β and IL-18 concentrations were quantified using the Mesoscale Discovery (MSD) U-PLEX Multiplex Kit (MSD, K15067M), following the manufacturer’s instructions. Samples were diluted 1:5 in kit diluent prior to analysis. The palmitate challenge assay was performed in three independent biological replicates. The concentration of IL-1β or IL-18 detected in the medium at each compound concentration was normalized to the value observed with DMSO control for data plotting.

The unbound fractions of TN-101 and TN-783 in palmitate-BSA matrix were determined by ultracentrifugation approach following the same procedures as used for *in vitro* determinations of plasma and brain binding, which are described in a following section. The protein-binding matrix consisted of palmitate–BSA in 150 mM NaCl (5 mM palmitate and 0.8 mM BSA; ∼6:1 molar ratio) that was diluted to 400 µM palmitate and 64 µM BSA with 150 mM NaCl, equilibrated at 37 °C for 10 min, and adjusted to pH 7.4. The IC_50_ values of the compounds in the THP-1 palmitate challenge assay were corrected for palmitate-BSA binding of the compounds using the following calculation: IC_50_ for IL-1β and IL-18 inhibition × unbound fraction of TN-101 or TN-783.

### Human and mouse whole blood assay

This protocol was adapted from (26) with some modification. Compounds were first serially diluted and dispensed into 384-well assay plates using a Tecan D300e liquid dispenser using a 9-point, 4-fold serial dilution. Upon receipt of fresh human or mouse blood, 190 µL was plated per well, into the plate with the compound dose response. After a 1-hour incubation at 37 °C, 10 uL lipopolysaccharide (LPS) (1 ng/mL final concentration) was added and incubated for 3.5 hours at 37 °C. 10 uL ATP (final concentration 5 mM) was added and incubated for 30 minutes at 37 °C followed by the addition of 250 uM menadione. The plates were spun at 2000 x g for 10 minutes at 4 °C after which 60 uL of the supernatant was used for detection of IL-1β. IL-1β was detected using the V-PLEX Human IL-1β or V-PLEX Mouse IL-1β kits (MSD) according to manufacturer’s instructions.

### Animals and diets

All procedures in animals were performed with adherence to ethical regulations and protocols approved by the Institutional Animal Care and Use Committee (IACUC) at the contract research facility. Male mice on a C57BL/6JGpt background were used for all experiments and were obtained from Gempharmatech. Mice were housed under a controlled temperature of 20-26°C and a 12-hour light cycle (lights on from 07:00 to 19:00) with *ad libitum* access to water and a standard chow (12.7% kcal from fat, 1010010, Jiangsu Xietong Pharmaceutical Bioengineering Co., Ltd.) or a high fat diet (60% kcal from fat, D12492, Research diet). DIO mice were maintained on the high fat diet for at least 15 weeks prior to study initiation. Animals were group-housed during the acclimation phase and subjected to mock dosing according to the intended route and frequency of study treatment, to habituate them to handling and procedures. Following acclimation, animals were randomized into treatment groups matched for baseline body weight and body composition, and transitioned to individual housing.

### Compound preparation for oral gavage dosing

TN-783 and TN-101 were prepared as a suspension in a 0.2% Tween 80 solution. Semaglutide was obtained from Novo Nordisk and formulated in saline. Dosing solutions were prepared on the first dosing day then weekly thereafter and were maintained at 4°C, on a laboratory rotator, until use. TN-783 and TN-101 suspension were remixed on dosing instances and administered via oral gavage and semaglutide was administered once daily via subcutaneous injection. Dosing volumes were standardized to 5 mL/kg body weight.

### Pharmacokinetic studies

To assess pharmacokinetics (PK), DIO and NC lean mice were dosed with TN-783 (50 mg/kg, b.i.d.) or TN-101 (1-10 mg/kg, b.i.d.) for 15 days. Blood samples were collected at the indicated timepoints after the first daily dose on day 15. Blood from TN-783-dosed animals was collected into EDTA-coated tubes supplemented with freshly prepared pefabloc (100 nM) at a 1:25 (v:v) ratio to inhibit soluble hydrolase activity. Samples were processed for plasma within half an hour of collection by centrifugation at approximately 4°C, 3200g for 10 minutes, then stored at -60°C or lower until LC-MS/MS analysis.

### Plasma and brain bioanalysis

Plasma and brain concentrations of TN-273 and TN-101 collected in PK and PK/PD studies were quantified using liquid chromatography-tandem mass spectrometry (LC-MS/MS). Brain samples were homogenized with 4 volumes (weight:volume) of phosphate buffered saline. Briefly, aliquots of 5-30 µl of calibration standards, quality control samples, blank matrix, and study samples were prepared for analysis with the addition of 100-600 µl (20x) of acetonitrile containing internal standard and mixing.

Precipitated contents were collected by centrifuging the samples, and the supernatants were removed for injection to the LC-MS/MS. Chromatographic separations were completed on an Acquity HSS T3 column (1.8 µm, 2.1 x 50 mm) with reversed phase gradient elution with mobile phases of 0.1% formic acid in water and acetonitrile. TN-273 and TN-101 were monitored with multiple-reaction monitoring with mass transitions of m/z 495.0 to 353.9 and m/z 459.1 to 357.9, respectively. Standard curve ranges were 0.002-6.0 µM for TN-273 and TN-101 in plasma and brain homogenates.

### *In vitro* determination of plasma and brain binding

The unbound fractions of TN-273 and TN-101 in mouse plasma and brain tissue were determined using an ultracentrifugation method. Test compounds were tested at 2 µM in mouse plasma containing Pefabloc (as an inhibitor of soluble hydrolases) and 1 µM in rat brain homogenate. Following equilibration for at least 10 min at 37°C, unbound compound was separated from bound compound by sedimentation of proteins using high centrifugal force (360,000 x g) for 2.5 h and quantified by LC-MS/MS. Concurrently, compounds were incubated at 37°C for 2.5 h in test matrices to verify the test article stability in matrix through the duration of the ultracentifugation procedures.

### Body weight and composition

Body weight and food intake was measured daily. Body composition was assessed by MRI (Minispec LF90, Bruker) at baseline and the indicated timepoints. At study termination, tissues were harvested and weighed, including white adipose depots (inguinal, epididymal, retroperitoneal, mesenteric) and representative muscles (gastrocnemius and tibialis anterior).

### Indirect calorimetry

Energy expenditure, food intake, and activity were measured using a Comprehensive Lab Animal Monitoring System (CLAMS, Columbus Instrument). Mice were housed in the CLAMS chambers for 4 consecutive days, with the first 48 hours used for acclimation. Data from the final 48 hours were averaged and presented as 24-hour hourly curves. Energy expenditure was calculated per animal and normalized to body weight. Respiratory exchange ratio (RER) was determined from volumes of oxygen consumption (VO_2_) and carbon dioxide production (VCO_2_).

### Oral glucose tolerance tests

For oral glucose tolerance test (OGTT), mice were subjected to daytime fasting from 10:00 for 6 hours. Glucose solution was administered orally (2 g/kg at a dosing volume of 5 ml/kg). Blood glucose was measured at baseline and at 15, 30, 60, and 120 minutes using a handheld glucometer (OneTouch VerioVue, Johnson & Johnson).

### Tissue collection

For all tissue collection, animals were deeply anesthetized with inhalation of 2-4% isoflurane. Following this, whole blood was collected via cardiac puncture into EDTA-coated tubes and centrifuged at 12,700 rpm for 7 min at 4°C before collecting the top plasma layer for analysis. Blood from TN-783-dosed animals was collected into EDTA-coated tubes supplemented with freshly prepared pefabloc (100 nM) at a 1:25 (v:v) ratio to inhibit soluble hydrolase activity. For studies in which hypothalami were collected for RNA sequencing or proteomics, mice were perfused with ice-cold PBS. The hypothalami were isolated and stored in -80°C until further processing. Inguinal and epididymal white adipose tissues from these animals were collected and drop-fixed in 4% paraformaldehyde and processed for paraffin embedding. For studies in which brains were processed for histology, mice were perfused with heparinized PBS (15,000 IU/L) followed by 10% neutral buffered formalin. Brains were collected, fixed, and paraffin-embedded.

### Histological assessment of hypothalamic inflammation

Coronal sections (4 µm) spanning the hypothalamus (bregma −1.4 to −3.0 mm) were collected and mounted on glass slides. Immunohistochemistry was performed on a Ventana Ultra platform (Roche Diagnostics) following deparaffinization and antigen retrieval (pH 8). Primary antibodies included anti-AgRP (R&D Systems AF634-SP), anti-Iba1 (Abcam ab178846), and anti-GFAP (Agilent/Dako Z0334), with appropriate anti-rabbit or anti-goat secondary antibodies. Slides were counterstained with DAPI and scanned on an Olympus VS200 microscope at 20X magnification with automated image stitching. Quantitative analysis focused on the arcuate nucleus (ARC), ventromedial hypothalamus (VMH), and dorsomedial hypothalamus (DMH). Regions of interest (ROIs) were manually delineated bilaterally, using AgRP immunoreactivity as a reference to reliably identify anatomical boundaries and ensure consistency across samples. For each animal, three coronal sections encompassing the ARC were analyzed (see Supplementary Figure 4 for anatomical reference). Fluorescent images were processed in ImageJ (77). Within each ROI, a fixed lower threshold was applied, and the number of suprathreshold pixels was measured. Pixel counts were normalized to the calibrated ROI area (mm²) to yield positive pixel density.

### Bulk RNA sequencing for hypothalamus

The hypothalamus was dissected from the brain of NC lean mice, vehicle-treated DIO mice, and TN-783-treated DIO mice following 29 d treatment with vehicle or TN-783 (n = 6 per group). To isolate RNA, each hypothalamus tissue sample was placed in a 1.5 mL centrifuge tube containing a 3 mm steel bead (Fisher, 50-206-7603). Subsequently, 350 µL of RLT Plus buffer (Qiagen) supplemented with β-mercaptoethanol was added to the tube. Homogenization was performed using a TissueLyser (Qiagen) for two cycles at 25 Hz for 3 minutes each. Following homogenization, samples were centrifuged at 16,000 × g for 3 minutes, and the clarified lysates were collected. RNA isolation was then carried out according to the manufacturer’s protocol using the RNeasy Plus Micro Kit (Qiagen, Cat. No. 74034). Purified RNA was transferred to 1.5 mL DNA LoBind tubes (Eppendorf), snap-frozen on dry ice, and stored at −80 °C until further processing for RNA sequencing.

For each of the 18 independent biological samples, a sequencing library was prepared with the QuantSeq 3’ mRNA-seq Library Prep Kit FWD for Illumina (Lexogen A01173). The UMI second strand synthesis module was used to introduce unique molecular identifiers (UMIs), following the protocol defined by the manufacturer. Briefly, oligo dT primers were hybridized to total RNA for reverse transcription, followed by RNA removal after first strand synthesis. UMIs were introduced during second strand synthesis and cDNA was purified using magnetic beads, followed by 18 cycles of PCR with dual indexes and PCR purification. Library quantity and quality were assessed with Qubit™ 1X dsDNA HS Assay Kits (Invitrogen Q33231) and Bioanalyzer High Sense DNA chip (Agilent 5067-4626). Libraries were pooled in equimolar ratios, and the sequencing pool was purified with magnetic beads to remove residual adaptor dimers. Sequencing was performed on an Illumina NovaSeq X Series 10B flowcell (100 bp single end) by SeqMatic (Fremont, CA, USA).

Data was processed using nf-core/rnaseq (v3.14.0) of the nf-core collection of workflows (78), using STAR (v2.7.9a) and salmon (v1.10.1) (79, 80) for alignment of reads to the GRCm39 mouse reference genome and quantification of Gencode (M31) gene models, respectively. UMIs were extracted from each aligned reads using the following regular expression: ^(?P.{6})(?P.{4}).*. The following extra arguments were provided to STAR to account for the 3’ tag nature of the library: --alignIntronMax 1000000 --alignIntronMin 20 --alignMatesGapMax 1000000 --alignSJoverhangMin 8 -- outFilterMismatchNmax 999 --outFilterType BySJout --outFilterMismatchNoverLmax 0.1 --clip3pAdapterSeq AAAAAAAA. Salmon quantification was performed without length correction by specifying the –noLengthCorrection argument. Differential expression analysis was performed with the edgeR (v4.6.2) and limma (v 3.64.1) Bioconductor packages using R v4.5.1. In brief, count matrices were subset to protein-coding genes, and library size factors were calculated with the TMM method. Gene-wise linear models with the experimental group as the single covariate were fit with the voomLmFit function, setting the sample weights argument to TRUE. Fold changes and p-values for the contrasts of interest were obtained with the eBayes function, and false discovery rates were calculated according to Benjamini, Hochberg, and Yekutieli. Volcano plots were generated with the ggplot2 R package (v3.5.2). Gene set enrichment analyses were performed with the fgsea R package (v1.34.0) with gene set collections from the msigdbr package (v24.1.0). Normalization of counts per million (cpm) was performed with the normLibSizes function from the edgeR R package, using the default TMM (trimmed mean of M-values) method by Robinson and Oshlack (2010). Principal component analysis revealed that two samples from the DIO_Vehicle group clustered away from all other samples. These two outliers exhibited exceptionally high expression of genes encoding prolactin and growth hormones, suggesting contamination with pituitary gland. Turning on the setting of sample weights automatically down-weighted the contribution of these samples. The top 25 genes altered in DIO relative to lean controls and rescued by TN-783 treatment were identified by overlapping the top 100 genes upregulated in DIO with the top 100 genes downregulated by TN-783, and vice versa, with ranking based on *p* values. Of the 26 overlapping genes, one with no characterized function was excluded. A heatmap of the remaining 25 genes was generated using normalized counts per million (cpm), which were first mean-normalized for each gene and then log2-transformed.

### Proteomics for hypothalamus

The hypothalamus was dissected from NC lean mice, vehicle-treated DIO mice, and TN-783-treated DIO mice following 14 d treatment with vehicle or TN-783 (n = 5 per group). After a single wash in ice-cold PBS, 600 µL ice-cold RIPA buffer containing 1× protease inhibitor, 1× phosphatase inhibitor, and 1× PMSF was added to each hypothalamic sample. Tissue homogenization was performed using a TissueLyser (Qiagen) for two cycles at 25 Hz for 1 min each. Homogenates were centrifuged at 12,000 × g for 10 min at 4 °C, and the supernatant was collected for downstream analysis. Protein concentration was determined using a BCA protein quantification kit. A total of 100 µg protein from each sample was subjected to digestion using the S-Trap™ micro column according to the manufacturer’s instructions, with the following modifications: 10 mM DTT and 30 mM IAA were used as reducing and alkylating reagents, respectively, in place of TCEP and MMTS; 50 mM TEAB, 0.2% FA in H₂O, and 75% ACN in 0.1% FA were used as elution solutions; and trypsin-digested peptides were dried using SpeedVac.

For LC-MS analysis, 200 ng peptides per sample were loaded onto a 25-cm C18 column and separated using a 60-min gradient on a NanoElute system (mobile phase A: 0.1% formic acid in H₂O; mobile phase B: 0.1% formic acid in acetonitrile). The separation gradient was as follows: 2–22% B from 0–37 min, 22–37% B from 37–46 min, 37–60% B from 46–50 min, 60–80% B from 50–52 min, followed by 80% B until the end. Eluted peptides were analyzed on a timsTOF Pro2 using DIA mode with the following settings: scan range, m/z 100–1700; 1/k₀ range, 0.7–1.35 V·s/cm²; ramp time, 100 ms; cycle time estimate, 1.38 s; collision energy, 20.00–59.00 eV.

Raw data were processed in DIA-NN. The mouse reference database was downloaded from UniProt (https://www.uniprot.org/). Search parameters were as follows: maximum missed cleavages, 1; maximum variable modifications, 2; fixed modification, carbamidomethylation at Cys; MS1 accuracy, 10 ppm; MS2 accuracy, 20 ppm. Differentially expressed protein (DEP) analysis was performed in R using the *limma* package. The top 25 proteins altered in DIO relative to lean controls and rescued by TN-783 treatment were identified by overlapping the top 100 proteins upregulated in DIO with the top 100 proteins downregulated by TN-783, and vice versa, ranked by *p* values. Of the 28 overlapping proteins, the top 25 were further ranked by fold change. A heatmap was generated from normalized protein abundance values, which were first mean-normalized for each protein and then log2-transformed.

### Statistical analysis

Data are reported as means ± SEM or as indicated in figures. Statistical analysis of data was performed in GraphPad Prism version 8 or later. Analysis of data from animal studies was performed using t test or one-way analysis of variance (ANOVA) with multiple comparison, or two-way ANOVA followed by multiple comparison test for data obtained from repeated measures of the same subjects.

