## SUPPLEMENTAL INFORMATION for "CNS-PENETRANT NLRP3 INHIBITOR ACHIEVES DURABLE WEIGHT LOSS AND REVERSES HYPOTHALAMIC INFLAMMATION IN DIET-INDUCED OBESITY"

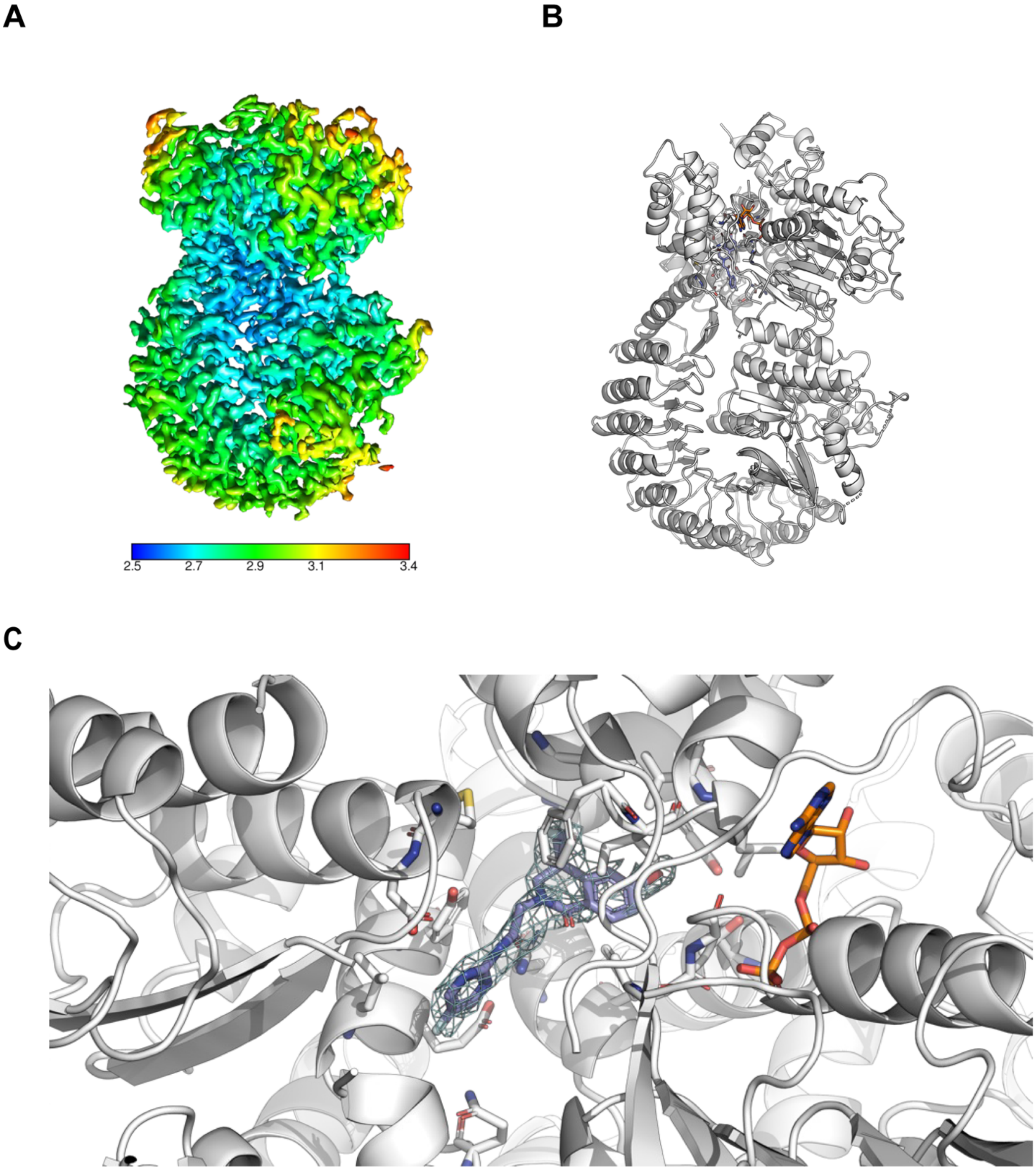


**Supplementary Figure 1 (related to Figure 1). Cryo-EM structure of NLRP3–NEK7 complex with bound NLRP3 inhibitor TN-551**

(**A**) CryoEM map of NLRP3 bound to NEK7 colored according to local resolution. (**B**) A reconstructed model of NLRP3 bound with NEK7 solved to 2.6 Å with clear density observed for TN-551 in the canonical inhibitor pocket (NLRP3 NACHT FP assay $IC_{50}$

values of 19.3 ± 4.5 nM) bound in a site adjacent to the ATP binding site (PDB: 9YJH). The interdomain cleft formed by the nucleotide-binding domain (NBD), helical domain 1 (HD1), winged helical domain (WHD), and HD2. TN-551 stabilizes the ADP-bound inactive state of NLRP3 and engages the same residues as MCC950 and NP3-253, a recently described pyridazine-based NLRP3 inhibitor, *via* the “intramolecular glue” mechanism previously described (1, 2). (**C**) TN-551 (blue) exhibits a distinct shape and interaction pattern from published complex structures. The central carbonyl of TN-551 forms a strong bifurcated hydrogen bond to both R578 and E629, while the dihydro-isoquinoline (DHIQ) bromo forms a halogen bond to Y443 and the F-pyrimidine interacts with S626 and E624. A structured water network further stabilizes the inactive NLR, with the TN-551 DHIQ core and central amide participating in solvent-mediated bridges to NLRP3. These features collectively explain the ability of TN-551 to lock NLRP3 in the inactivated state.

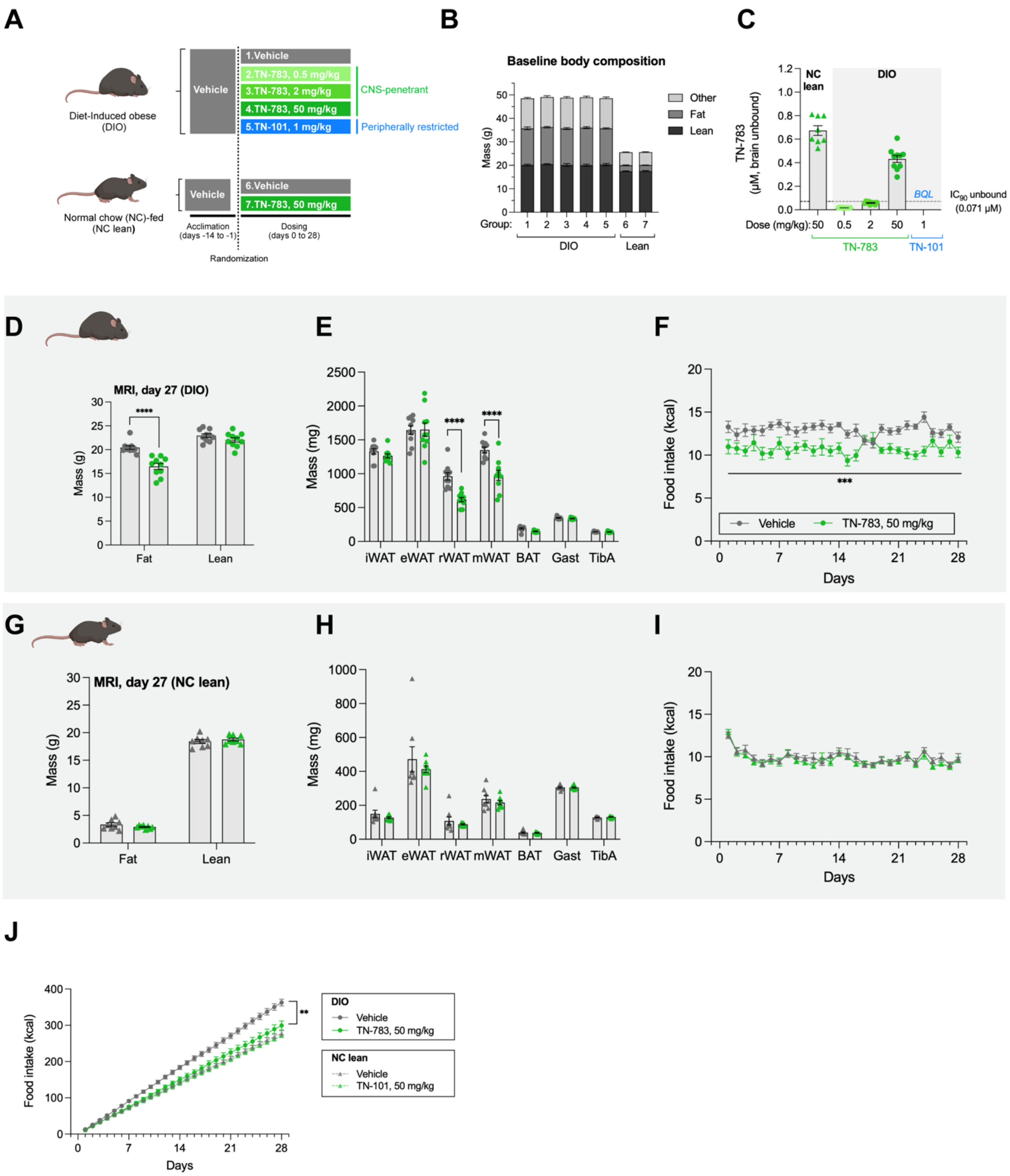


**Supplementary Figure 2 (related to Figure 3). CNS-penetrant NLRP3 inhibitor promoted reduced fat mass in association with lowered food intake.**

(**A**) Schematic of compound dosing in DIO and NC lean mice. (**B**) Baseline body composition in DIO and lean mice. (**C**) TN-783 and TN-101 brain exposure. (**D-F**) Fat and lean masses, weight of representative adipose tissues and muscles, and food intake in TN-783-treated DIO mice. (**G-I**) Fat and lean masses, weight of representative adipose tissues and muscles, and food intake in TN-783-treated NC lean controls. (**J**) Cumulative food intake in DIO and NC lean mice. Data are mean ± SEM of N=8-10/group. Statistical significance for TN-783 effect was determined by a two-way ANOVA (**E, H, G**) followed by Šídák's multiple comparison tests against vehicle-treated controls (**D-E**, **G-H**) (**, $p<0.01$, ***, $p<0.001$, ****, $p<0.0001$). iWAT: inguinal white adipose tissue, eWAT: epididymal white adipose tissue, rWAT: retroperitoneal white adipose tissue, mWAT: mesenteric white adipose tissue, BAT: brown adipose tissue, Gast: gastrocnemius muscle, TibA: tibialis anterior muscle.

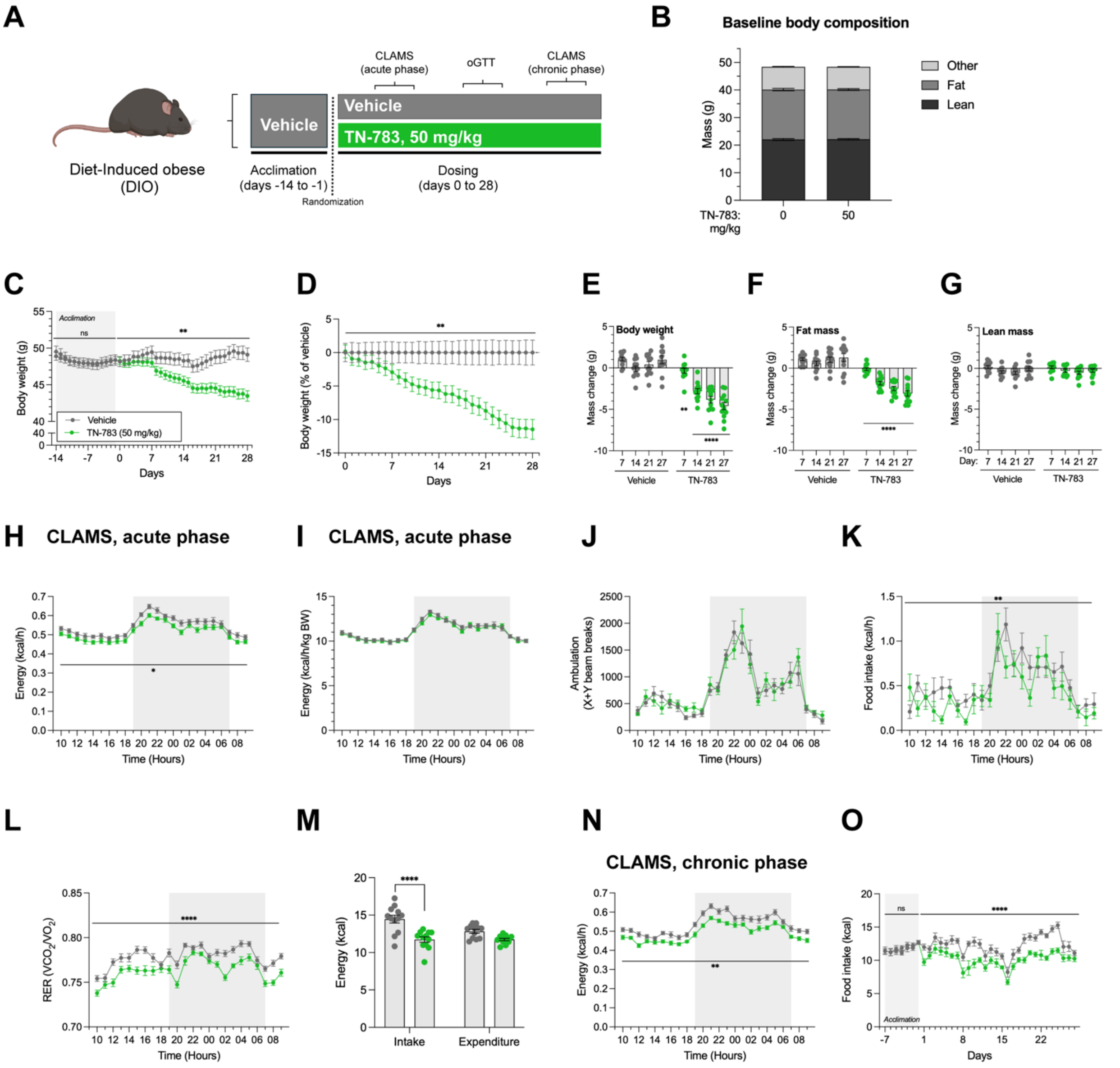


**Supplementary Figure 3 (related to Figure 3). TN-783 induced weight loss was characterized by fat mass reduction and driven by reduction in food intake.** (**A**) Schematic of TN-783 dosing in DIO mice. Animals underwent CLAMS measurements at room temperature in the acute (days 7-11) and chronic (days 21-25) phases of TN-783 dosing. OGTT was performed after 15 days of TN-783 dosing. (**B**) Baseline body composition in DIO mice. (**C-D**) Absolute and vehicle-adjusted body weight of DIO mice treated with TN-783. (**E-G**) Weekly changes in body weight, fat and lean masses. (**H-L**) Absolute and body weight-normalized energy expenditure, physical activity, food intake, and RER over 24 hours in the acute phase of TN-783 dosing. Grey areas indicate dark/lights-off period. (**M**) Total energy intake and energy expenditure over the 24-hour period measured in the acute phase of the study. (**N**) Absolute energy expenditure over 24 hours in the chronic phase of TN-783 dosing. (**O**) Daily food intake over the acclimation and dosing periods. Data are mean ± SEM of N=12/group. Statistical significance for TN-783 effect was determined by a two-way ANOVA (**C-D**, **H-L, N-O**) followed by Šídák's multiple comparison tests against vehicle-treated controls (**E-G, M**) (*, $p<0.05$. **, $p<0.01$, ****, $p<0.0001$).

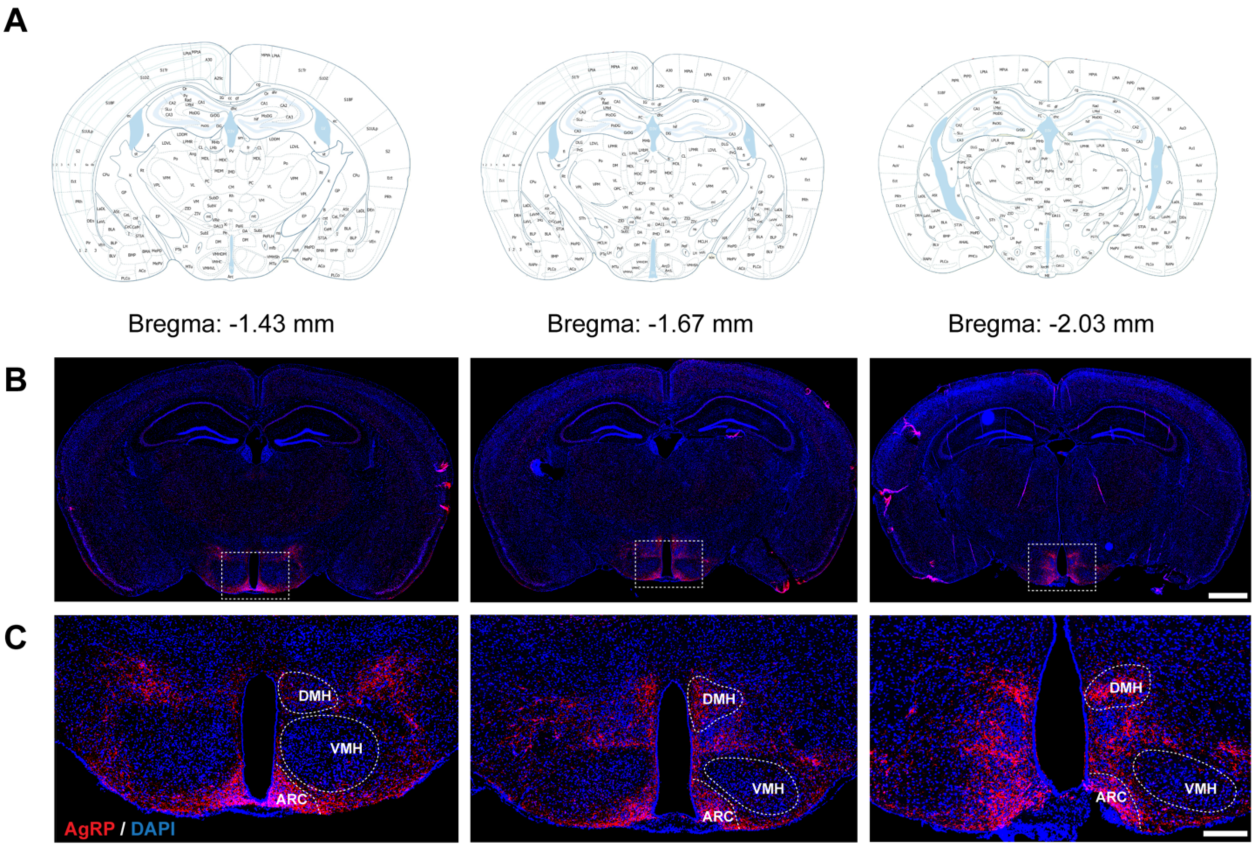


**Supplementary Figure 4 (related to Figure 4). Identification of distinct hypothalamic nuclei through AgRP immunostaining.**
(**A**) Illustrations of stereotaxic coordinates at –1.43 mm, –1.67 mm, and –2.03 mm from bregma, used to anatomically identify regions of interest. (**B**) Representative coronal sections at the corresponding bregma levels showing AgRP immunofluorescence (red) and DAPI nuclear staining (blue). Dashed boxes highlight the hypothalamic regions selected for further analysis. (**C**) Higher-magnification views of the hypothalamus with outlines of the arcuate nucleus (ARC), ventromedial hypothalamus (VMH), and dorsomedial hypothalamus (DMH). AgRP staining was used to confirm regional boundaries for downstream quantification. Scale bar: (B) 500 µm, (C) 100 µm.

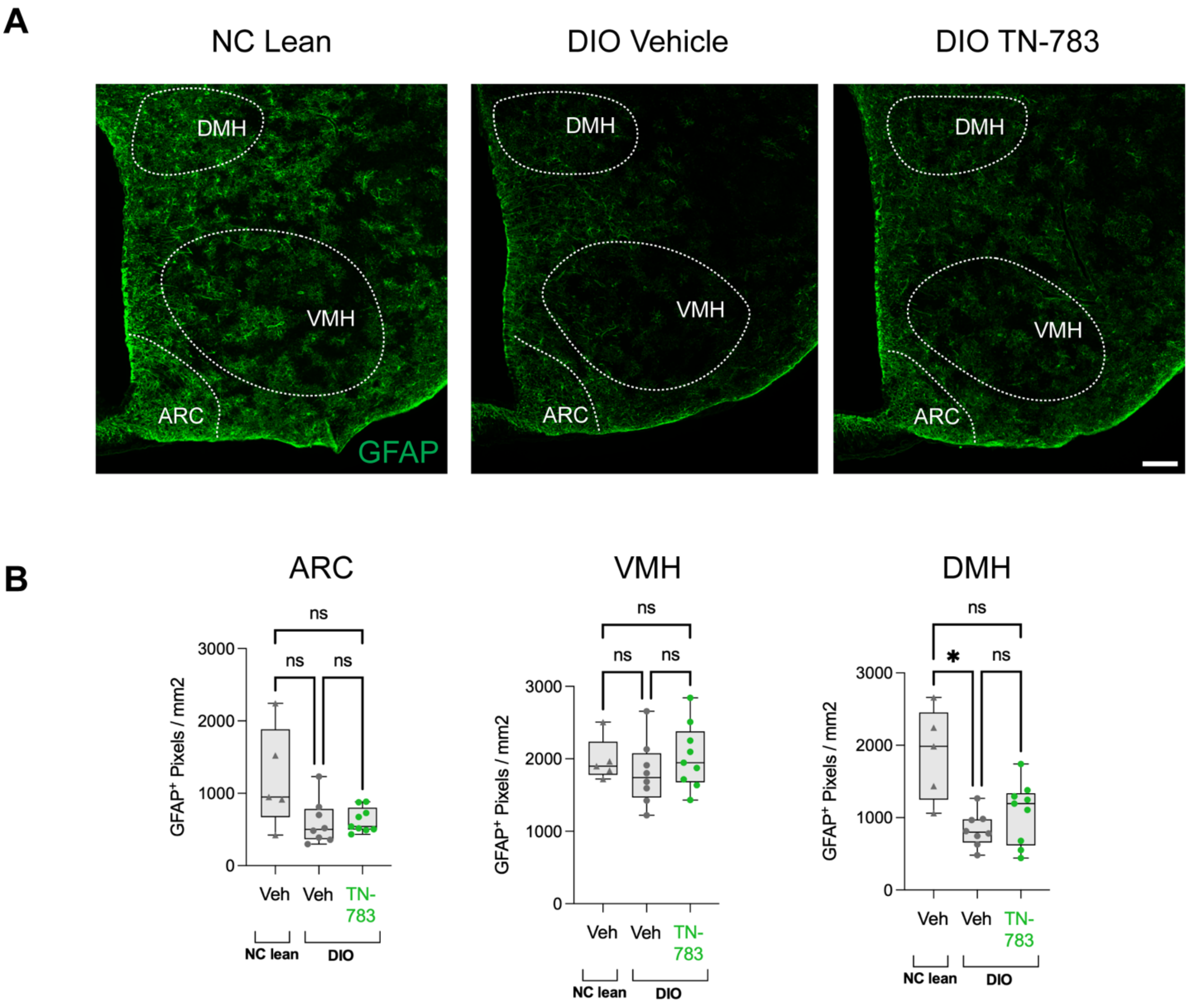


**Supplementary Figure 5 (related to Figure 4). GFAP quantification in the ARC, VMH and DMH.**

Representative sections showing GFAP immunoreactivity in the hypothalamus of vehicle-treated NC lean control, vehicle- and TN-783-treated DIO animals. Scale bar: 100 µm. ARC: arcuate nucleus, VMH: ventromedial hypothalamus, DMH: dorsomedial hypothalamus (**B**) Quantification of GFAP immunostaining. Data are shown as median with interquartile range (IQR); whiskers indicate min–max. Statistical significance for TN-783 effect was determined by a one-way ANOVA followed by Dunnett's multiple comparison tests against vehicle-treated DIO controls (*, $p<0.05$. **).

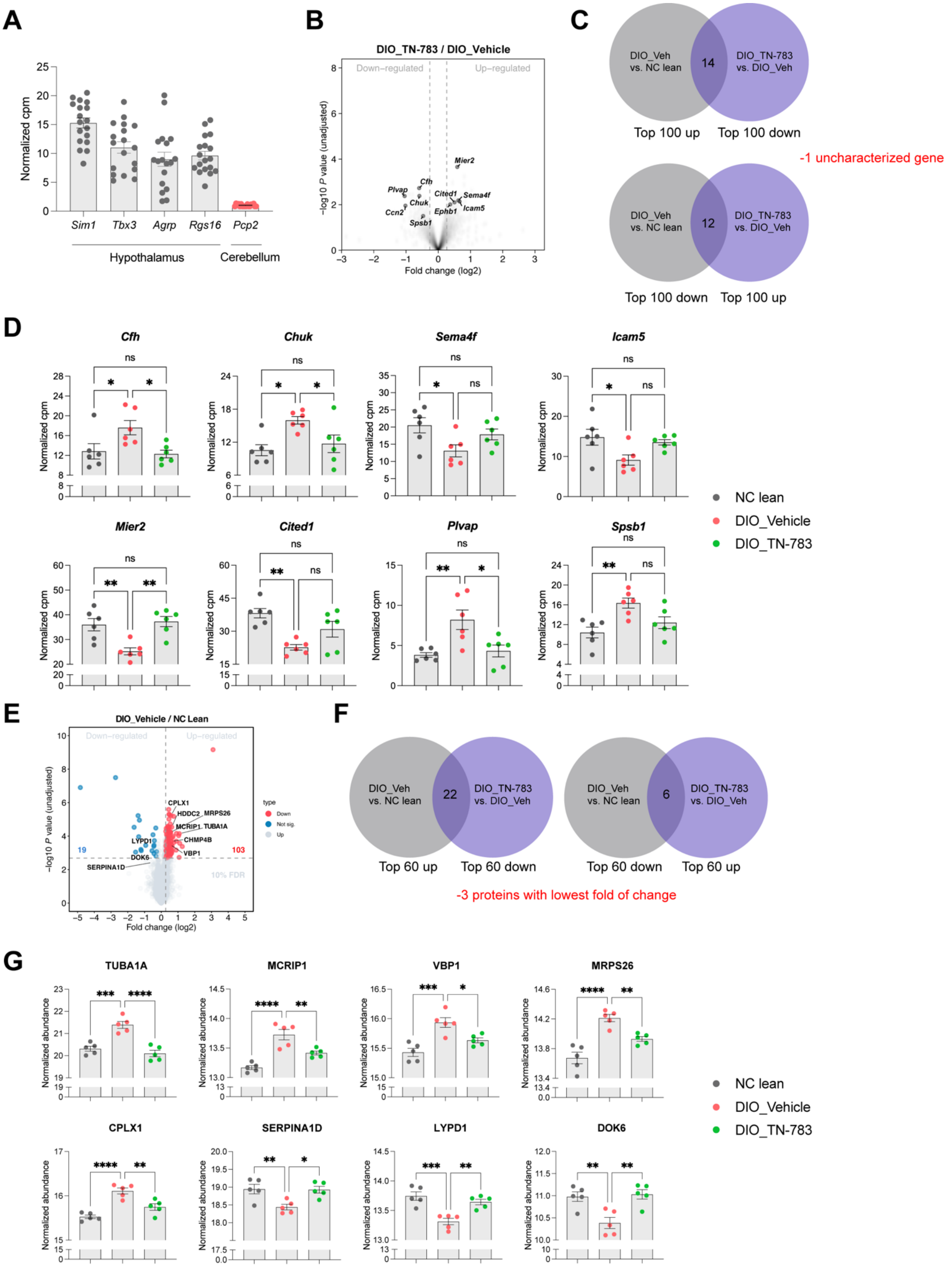
A
Normalized cpm
Sim1
Tbx3
Agrp
Rgs16
Pcp2
Hypothalamus
Cerebellum
B
DIO_TN-783 / DIO_Vehicle
Down–regulated
Up–regulated
–log10 P value (unadjusted)
Fold change (log2)
Mier2
Cfh
Plvap
Chuk
Ccn2
Spsb1
Cited1
Sema4f
Ephb1
Icam5
C
DIO_Veh vs. NC lean
DIO_TN-783 vs. DIO_Veh
14
Top 100 up
Top 100 down
-1 uncharacterized gene
12
Top 100 down
Top 100 up
D
Cfh
Chuk
Sema4f
Icam5
Mier2
Cited1
Plvap
Spsb1
Normalized cpm
ns
NC lean
DIO_Vehicle
DIO_TN-783
E
DIO_Vehicle / NC Lean
Down–regulated
Up–regulated
–log10 P value (unadjusted)
Fold change (log2)
type
Down
Not sig.
Up
19
103
10% FDR
CPLX1
HDDC2
MRPS26
MCRIP1
TUBA1A
CHMP4B
VBP1
LYPD1
DOK6
SERPINA1D
F
DIO_Veh vs. NC lean
DIO_TN-783 vs. DIO_Veh
22
Top 60 up
Top 60 down
6
Top 60 down
Top 60 up
-3 proteins with lowest fold of change
G
TUBA1A
MCRIP1
VBP1
MRPS26
CPLX1
SERPINA1D
LYPD1
DOK6
Normalized abundance
NC lean
DIO_Vehicle
DIO_TN-783

**Supplementary Figure 6 (related to Figure 4). TN-783 partially reversed DIO-induced transcriptomic and proteomic changes in hypothalamus**

(**A**) Normalized counts per million (cpm) for the four hypothalamic marker genes (Sim1, *Tbx3*, *Agrp* and *Rgs16*) and a control gene specific for cerebellum (*Pcp2*) detected in the hypothalamic samples used for RNA sequencing. Each dot represents value from one mouse. (**B**) Volcano plot for transcriptomic changes in the hypothalamus of TN-783-treated DIO mice relative to vehicle-treated DIO mice. Note that none of the changes reached FDR < 10% but the labeled genes are examples of genes altered by TN-783 in the opposite direction of DIO-induced changes (compare with **Figure 4C**). (**C**) Venn diagrams illustrating the selection of top 25 genes rescued by TN-783 treatment in the hypothalamus of DIO mice based on RNA sequencing. (**D**) Normalized cpm (mean ± SEM) for 8 genes showing significant DIO-induced changes rendered insignificant with TN-783 treatment. Statistical significance was demonstrated by one-way ANOVA followed by Tukey's multiple comparison tests for all 3 pairs of comparisons (*, p<0.05. **, p<0.01. ns, non-significant). (**E**) Volcano plots showing proteomic changes in the hypothalamus of DIO mice relative to NC lean controls. (**F**) Venn diagrams illustrating the selection of top 25 proteins rescued by TN-783 in the hypothalamus of DIO mice based on proteomics. (**G**) Normalized abundance (mean ± SEM) for 8 proteins showing significant DIO-induced changes with significant reversal by TN-783 treatment. Statistical significance was demonstrated by one-way ANOVA followed by Dunnett's multiple comparison tests against vehicle-treated DIO (*, p<0.05. **, p<0.01.***, p<0.001. ***, p<0.0001).

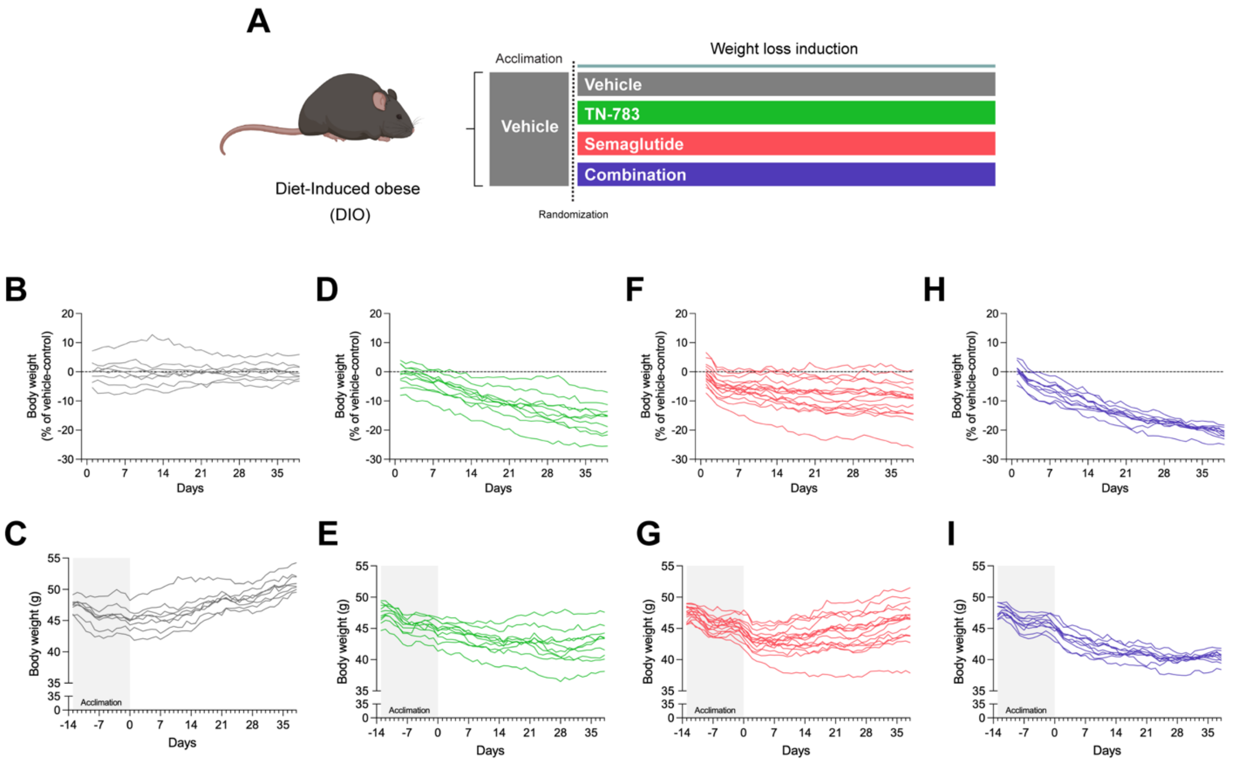


**Supplementary Figure 7 (related to Figure 6). TN-783 potentiated and improved uniformity in response to semaglutide-induced weight loss.**
(**A**) Study schematic of TN-783 and semaglutide dosing in diet-induced obese (DIO) mice. (**B-I**) Body weight (vehicle-adjusted and absolute) in DIO mice treated with vehicle, TN-783, semaglutide, or their combination during the weight loss induction period. Data are shown as trajectories of individual animals with each line representing one animal, N=8-16/group.

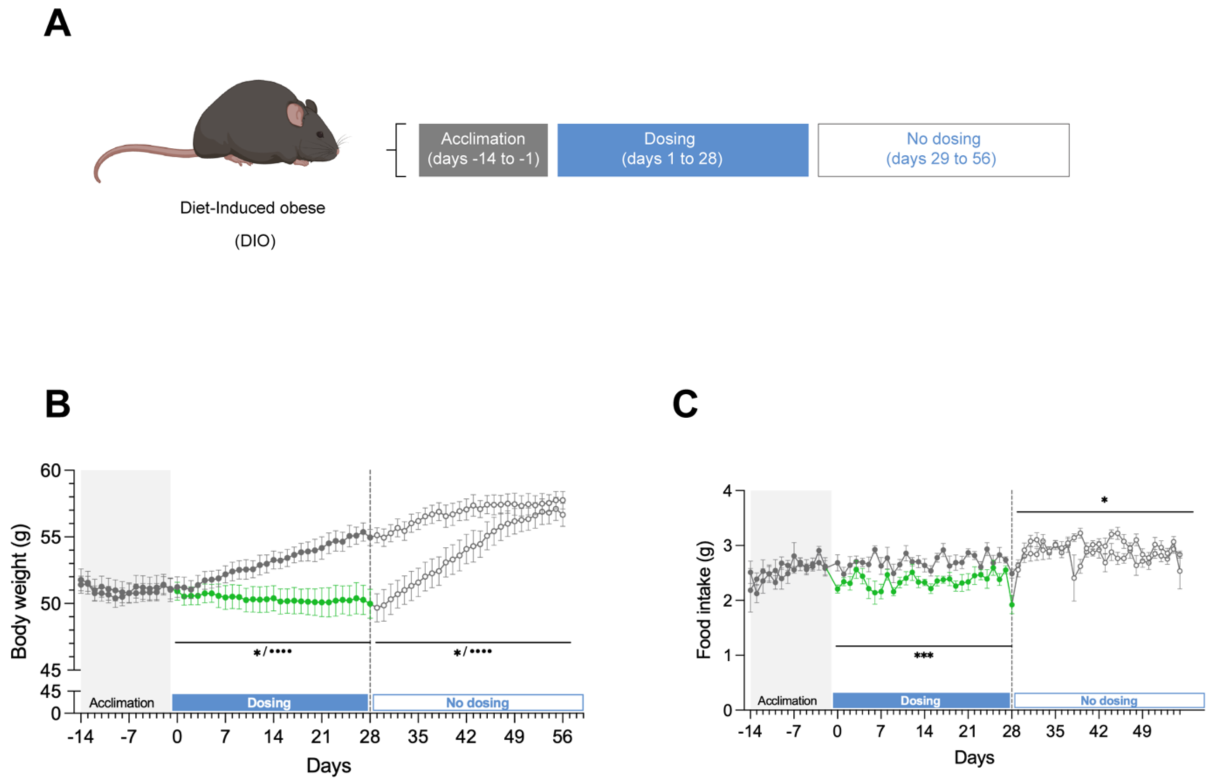


**Supplementary Figure 8 (related to Figure 6). TN-783 effect in suppressing body weight and feeding in DIO mice was reversible upon treatment cessation.**
(**A**) Study schematic of TN-783 dosing in diet-induced obese (DIO) mice. (**B**) Absolute body weight. (**C**) Food intake. Data are mean ± SEM of N=7-8/group. Statistical significance for TN-783 effect was determined by a two-way ANOVA against vehicle-treated DIO. Asterisks (*) indicate a significant main effect of drug, whereas bullet point (•) indicate a significant drug x time interaction (*, $p<0.05$, ***, $p<0.001$, ****/••••, $p<0.0001$).

**Supplementary Table 1.** Cryo-EM data collection, refinement, and validation statistics for the NLRP3–NEK7–TN-551 complex.

| **NEK7&NLRP3 complex** | (**PDB ID:** 9YJH )<br>(**EMDB ID:** EMD-73026) |
|---|---|
| **Data collection and Processing (for each dataset)** | |
| Microscope | Titan Krios |
| Voltage (kV) | 300 |
| Camera | Gatan K3 Summit |
| Magnification | 105,000 |
| Pixel size at detector (Å/pixel) | 0.83 |
| Total electron exposure ($e^-$/Å$^2$) | 80.5 |
| Number of frames collected during exposure | 50 |
| Defocus range (μm) | -1.0 ~ -2.5 |
| Phase plate (if used) | N/A |
| - phase shift range (in degrees) | N/A |
| - number of images per phase plate position | N/A |
| Automation software | SerialEM |
| Tilt angle | 0 |
| Energy filter slit width (eV) | 20 |
| Micrographs collected (no.) | 5562 |
| Micrographs used (no.) | 4039 |
| Total extracted particles (no.) | 3,591,811 |
| **For each reconstruction:** | |
| Monomer Refined particles (no.) | 340,724 |
| Monomer Final particles (no.) | 340,724 |
| Monomer Resolution (FSC 0.143, Å) | 2.61 |
| Monomer Resolution range (local, Å) | 2.0-3.2 |
| Monomer map sharpening B factor (Å$^2$) | 55.6 |
| Point-group or helical symmetry parameters | C1 |
| Map sharpening methods | cryoSPARC v3.0.1; |
| **Monomer Model composition** | |
| Protein | 1026 |
| Ligands | 4 |
| **Monomer Model Refinement** | |
| Refinement package | CCPEM 1.5.0 |
| Chains | 3 |
| Atoms | 16,627 (Hydrogens: 8,345) |
| Protein residues | Protein: 1026 Nucleotide:0 |
| Ligands | MG:2 |
| | DN1:1 |
| | ADP:2 |
| R.m.s. deviations from ideal values | |
| Bond lengths (Å) | 0.003 (0) |
| Bond angles (°) | 0.543 (6) |
| **Monomer Validation** | |
| MolProbity score | 1.18 |
| CaBLAM outliers | 0.82 |
| Clashscore | 1.63 |
| Rotamers outliers (%) | 1.74 |
| C-beta deviations | 0.00 |
| Ramachandran plot | |
| Favored (%) | 97.60 |
| Outliers (%) | 0 |

**Supplementary Table 2 (related to Figure 1).** Summary of *in vitro* potency of TN-101 and TN-783 across human and mouse NLRP3-dependent assays.

| **Assay** | **TN-101 $IC_{50}$ (nM)** | **TN-783 $IC_{50}$ (nM)** |
|---|---|---|
| Human NLRP3 fluorescent probe displacement (FP) | 29.8 ± 4.9 | 19.3 ± 2.2 |
| LPS/Nigericin stimulated THP-1 (IL-1β) | 5.58 ± 0.47 | 45.9 ± 3.0 |
| Palmitate-stimulated THP-1 (IL-1β) | 5.97 ± 0.94* | 5.17 ± 0.73* |
| Palmitate-stimulated THP-1 (IL-18) | 12.2 ± 4.3* | 12.3 ± 1.4* |
| LPS/ATP stimulated mouse whole blood (IL-1β) | 26.6 ± 7.1 | 174 ± 37 |
| LPS/ATP stimulated human whole blood (IL-1β) | 3.39 ± 0.83 | 23.1 ± 6.0 |

Calculated $IC_{50}$ values from three independent experiments are summarized as mean ± SEM.

*$IC_{50}$ values were corrected for palmitate-BSA binding (fraction unbound: 0.347 for TN-101, 0.152 for TN-783).

**Supplementary Table 3 (related to Figure 2).** Steady-state unbound plasma and brain exposures of TN-783 and TN-101 in DIO mice relative to *in vitro* potency.

| | Dose (mg/kg/dose, BID) | Observed steady-state unbound plasma exposure (μM) | | Predicted steady-state unbound brain exposure (μM) | | Fold of unbound $IC_{90}$ in plasma | | Fold of unbound $IC_{90}$ in brain | |
|---|---|---|---|---|---|---|---|---|---|
| | | Peak | Trough | Peak | Trough | Peak | Trough | Peak | Trough |
| **TN-783** | 50 | 0.91 | 0.60 | 0.37 | 0.25 | 13x | 8.5x | 5.3x | 3.5x |
| **TN-101** | 1 | 0.16 | 0.027 | 0.003 | 0.0004 | 7.0x | 1.2x | 0.1x | <0.1x |
| | 3 | 0.63 | 0.11 | 0.010 | 0.0018 | 27x | 4.8x | 0.4x | 0.1x |
| | 10 | 1.26 | 0.28 | 0.020 | 0.0045 | 55x | 12x | 0.9x | 0.2x |

DIO mice were dosed twice daily (*b.i.d.*) for 15 days with TN-783 (50 mg/kg/dose) or TN-101 (1, 3, or 10 mg/kg/dose). Steady-state unbound plasma concentrations (peak and trough) were measured, and steady-state unbound brain concentrations were predicted from plasma concentrations by applying the measured unbound brain-to-plasma partition coefficients ($K_{p,uu}$) obtained from separate PK studies (TN-783: 0.41, TN-101: 0.016,). Peak values correspond to 1-2 hours post-first daily dose on Day 15, and trough values correspond to 24 hours post-first daily dose on Day 15. Fold of $IC_{90}$ values for plasma and brain were calculated relative to the in vitro $IC_{90}$ of IL-1β secretion determined with LPS/ATP stimulated mouse whole blood corrected for plasma protein binding.

TN-783 $IC_{90,u}$ was derived from the mouse whole blood assay ($IC_{50}$ = 174 ± 37 nM; assuming a Hill slope of 1, corresponding to an $IC_{90}$ of ~1566 nM), and after correction for plasma protein binding (fu = 0.045), was calculated to be 71 nM. TN-101 $IC_{90,u}$ was similarly derived from the mouse whole blood assay ($IC_{50}$ = 26.6 ± 7.1 nM; assuming a Hill slope of 1, corresponding to an $IC_{90}$ of ~239.4 nM), and after correction for plasma protein binding (fu = 0.110), was calculated to be 26.3 nM.

**Supplementary Table 4 (related to Figure 4):** Hallmark pathways significantly altered in the hypothalamus of DIO_Vehicle mice relative to that of NC lean mice (FDR < 0.1).

| Gene set | N* | Unadjusted p value | FDR |
|---|---|---|---|
| HALLMARK_OXIDATIVE_PHOSPHORYLATION | 189 | 1.06E-21 | 5.30E-20 |
| HALLMARK_ADIPOGENESIS | 178 | 1.33E-08 | 3.33E-07 |
| HALLMARK_FATTY_ACID_METABOLISM | 127 | 9.38E-07 | 1.56E-05 |
| HALLMARK_TNFA_SIGNALING_VIA_NFKB | 118 | 0.000224128 | 0.00280161 |
| HALLMARK_BILE_ACID_METABOLISM | 73 | 0.001989816 | 0.01943281 |
| HALLMARK_REACTIVE_OXYGEN_SPECIES_PATHWAY | 43 | 0.002716742 | 0.01943281 |
| HALLMARK_INTERFERON_GAMMA_RESPONSE | 100 | 0.002720593 | 0.01943281 |
| HALLMARK_COAGULATION | 62 | 0.003710354 | 0.02318971 |
| HALLMARK_ANDROGEN_RESPONSE | 85 | 0.006561075 | 0.03155463 |
| HALLMARK_XENOBIOTIC_METABOLISM | 123 | 0.00673667 | 0.03155463 |
| HALLMARK_INFLAMMATORY_RESPONSE | 85 | 0.007564153 | 0.03155463 |
| HALLMARK_MYC_TARGETS_V1 | 182 | 0.007573111 | 0.03155463 |
| HALLMARK_COMPLEMENT | 113 | 0.008761093 | 0.03369651 |
| HALLMARK_HYPOXIA | 145 | 0.013612815 | 0.0464682 |
| HALLMARK_G2M_CHECKPOINT | 131 | 0.014354305 | 0.0464682 |
| HALLMARK_MITOTIC_SPINDLE | 160 | 0.014869824 | 0.0464682 |
| HALLMARK_P53_PATHWAY | 151 | 0.017925987 | 0.05272349 |
| HALLMARK_HEDGEHOG_SIGNALING | 31 | 0.021395533 | 0.05943204 |
| HALLMARK_DNA_REPAIR | 131 | 0.030386502 | 0.07996448 |
| HALLMARK_ESTROGEN_RESPONSE_LATE | 126 | 0.032672557 | 0.08168139 |

*Number of genes in each gene set
